# Structure, biosynthesis, and bioactivity of nostolysamides

**DOI:** 10.64898/2026.01.31.703028

**Authors:** Enleyona Weir, Ivan Anterola, Wilfred A. van der Donk

## Abstract

A recent genome mining study identified class II lanthipeptides encoded in *Nostoc punctiforme* PCC 73102 that contain acyl groups conjugated to Lys side chains. The structure and bioactivity of these peptides, named nostolysamides, were not determined. In this study, we produced the nostolysamides by co-expression of the NpuA precursor peptide with an N-terminal SUMO tag with the class II lanthipeptide synthetase NpuM in *Escherichia coli.* All four lanthionine and methyllanthionine residues were shown to have the DL configuration by Marfey’s analysis. Tandem mass spectrometry and mutagenesis studies indicate an N-terminal non-overlapping methyllanthionine ring and three overlapping rings at the C-terminus for which the most likely ring pattern is proposed. The NpuM lanthipeptide synthetase is a member of the ProcM-clade and catalyzes ring formation with both C-to-N and N-to-C directionality. After removal of the leader peptide, the resulting lanthipeptide exhibits antibacterial as well as antifungal activity against *Candida* species by disrupting cell membranes. Antibacterial activity is shown not to involve lipid II. The biosynthetic gene cluster also encodes an acetyltransferase NpuN that transfers long chain acyl groups to the side chain of a Lys residue in position 1 of the precursor peptide. *In vitro* studies of NpuN show relatively broad substrate specificity with NpuN conjugating various acyl groups from acyl-CoA substrates to Lys1 in the nostolysamides. The acylation did not appreciably change the antifungal and antimicrobial activity of nostolysamide showing that it is not required for these activities.

## Introduction

Invasive fungal infections have been estimated to affect over 6.5 million people worldwide each year, resulting in 3.8 million deaths.^1^ This figure represents a significant increase over previous estimates, highlighting a growing global public health concern related to fungal infections and antimicrobial resistance. Candidiasis accounts for 24% of invasive fungal infections, making it a significant contributor.^1^ *Candida albicans*, responsible for most *Candida* infections, is a commensal yeast that inhabits the skin and various mucosal surfaces of the human body.^2, 3^ *Candida* species are opportunistic pathogens, especially in immunocompromised patients, and can cause various diseases with high mortality rates.^1, 4^

The rise of antifungal resistance poses a significant threat in the clinical setting. Therefore, efforts have been made to identify and characterize novel molecules with antifungal activity. Natural products constitute an important class of antifungal compounds. Amongst natural products, ribosomally synthesized and post-translationally modified peptides (RiPPs) are a rapidly growing family of molecules with a wide spectrum of biological activities, including antifungal activities.^5^ Lanthipeptides are one of the largest families of RiPPs based on current genomes.^6, 7^ They are characterized by the presence of a thioether crosslink formed from a Ser/Thr and a Cys residue.^8^ Lanthipeptides are grouped into five classes based on their biosynthetic enzymes and the mechanisms by which they form the thioether linkages. The biosynthesis of class I lantibiotics, such as the widely used food preservative nisin, involves a dehydratase (LanB) and a cyclase (LanC) to form the crosslinks.^9^ In contrast, class II peptides are modified by a single bifunctional enzyme (LanM) responsible for dehydration and cyclization.^8^ The dehydration domain of LanM enzymes does not have sequence or structural homology with LanB proteins, but the cyclization domains of LanMs do have homology with the LanC proteins.^10–12^ Class III and IV lanthipeptides employ trifunctional enzymes (LanKC-type and LanL, respectively).^13, 14^ Their dehydration domains are unrelated to LanB and LanM enzymes, but their cyclization domains have sequence and structural homology with the LanC cyclases.^15–17^ The recently identified class V lanthipeptides require unique enzymes, distinct from those in other classes, to mediate the post-translational modifications.^18–22^

Pinensins A and B are the first and only example of an antifungal lanthipeptide, which were isolated from *Chitinophaga*.^23^ The pinensins constitute a two-component class I lanthipeptide. In this study, we characterize another lanthipeptide that exhibits activity against *Candida* species that is a member of class II. During the biosynthesis of class II lanthipeptides, a bifunctional enzyme (LanM) carries out dehydration of Ser and Thr residues to their corresponding dehydroamino acids dehydroalanine (Dha) and dehydrobutyrine (Dhb), and cyclization by Michael-type addition of Cys residues to the Dha/Dhb residues.^24^ These post-translational modifications are performed on a precursor peptide (LanA) to form lanthionine (Lan from Ser/Cys) or methyllanthionine (MeLan from Thr/Cys) rings.^8^ The modifications are typically introduced in the C-terminal part of the precursor peptide, LanA, called the core peptide, whereas an N-terminal leader peptide is removed by a peptidase to generate the mature natural product.^8^ In some instances, additional modifications besides (methyl)lanthionine rings are introduced by enzymes encoded in the biosynthetic gene cluster (BGC).^25^ For instance, general control non-repressible 5 (GCN5)-related N-acetyltransferases (GNATs) attach acyl groups.^26, 27^ Acylation of peptides with lipid chains typically increases stability,^28^ potency, or metabolic half-life.^28–32^ Only a few known lanthipeptides are lipidated by GNATs, such as microvionin, the first example of a class III lanthipeptide that contains an N-terminal lipid.^33^ Other N-terminally lipidated class III lanthipeptides include the goadvionins^34^ and the lipoavitides.^35, 36^ Piel and Vagstad and colleagues discovered lanthipeptides in which not the N-terminus but the side chains of ornithines are lipidated.^37^ They termed these RiPPs in which GNATs lipidate amino acid side chains selidamides, and characterized two members, kamptornamide and phaeornamide, encoded in cyanobacterial and marine α-proteobacterial BGCs, respectively.^37^ The authors also showed that lanthipeptide BGCs encoding these GNATs are widespread, and for one BGC from *Nostoc punctiforme* PCC 73102 (*npu*), they demonstrated acylation of the side chain of Lys in position 1.^37^ The structure and bioactivity of this lanthipeptide could not be determined because of the low production level. Independently, as part of a program to produce RiPPs using a Fast, Automated, Scalable, high-Throughput pipeline for RiPP discovery (FAST-RiPPs),^38^ our group had targeted the same BGC. Here, we report further characterization of the product of the *npu* cluster and its bioactivity.

## Result and Discussion

### Heterologous co-expression of the precursor peptide NpuA with NpuM

A previous bioinformatics genome-mining study mapped lanthipeptide diversity across sequenced genomes, identifying thousands of uncharacterized peptides with unique precursor peptide sequences that would yield new ring patterns.^39^ Within the class II lanthipeptides reported in this bioinformatic study, a group of lanthipeptide precursors was identified in cyanobacteria that includes the *npu* BGC in *Nostoc punctiforme* PCC 73102 (**Fig. 1A**). As noted previously,^37^ this BGC encodes a precursor peptide NpuA (WP_012412980.1), a class II lanthionine synthetase NpuM (WP_012412979.1), and a GNAT NpuN (WP_012412981.1) (**Fig. 1B**). To explore the distribution of *npu* BGCs, we also performed PSI-BLAST searches of the NCBI non-redundant database using the precursor peptide NpuA as query, yielding several BGCs that encode homologs of all three proteins (**Fig. 1C, D**). NpuA consists of a leader peptide (LP) of 78 amino acid residues, which is required for substrate recognition but does not undergo post-translational modification. This LP is a member of the Nif11 leader peptide family^42, 43^ and has a predicted Gly-Gly protease cleavage site, which is anticipated to be removed by the C39 peptidase domain of NpuT, a peptidase-containing ATP-binding cassette transporter (PCAT).^44, 45^ The core peptide of NpuA is composed of 25 amino acids, including two Thr residues, three Ser residues, and four Cys residues (**Fig. 1B**).

**Figure 1.**
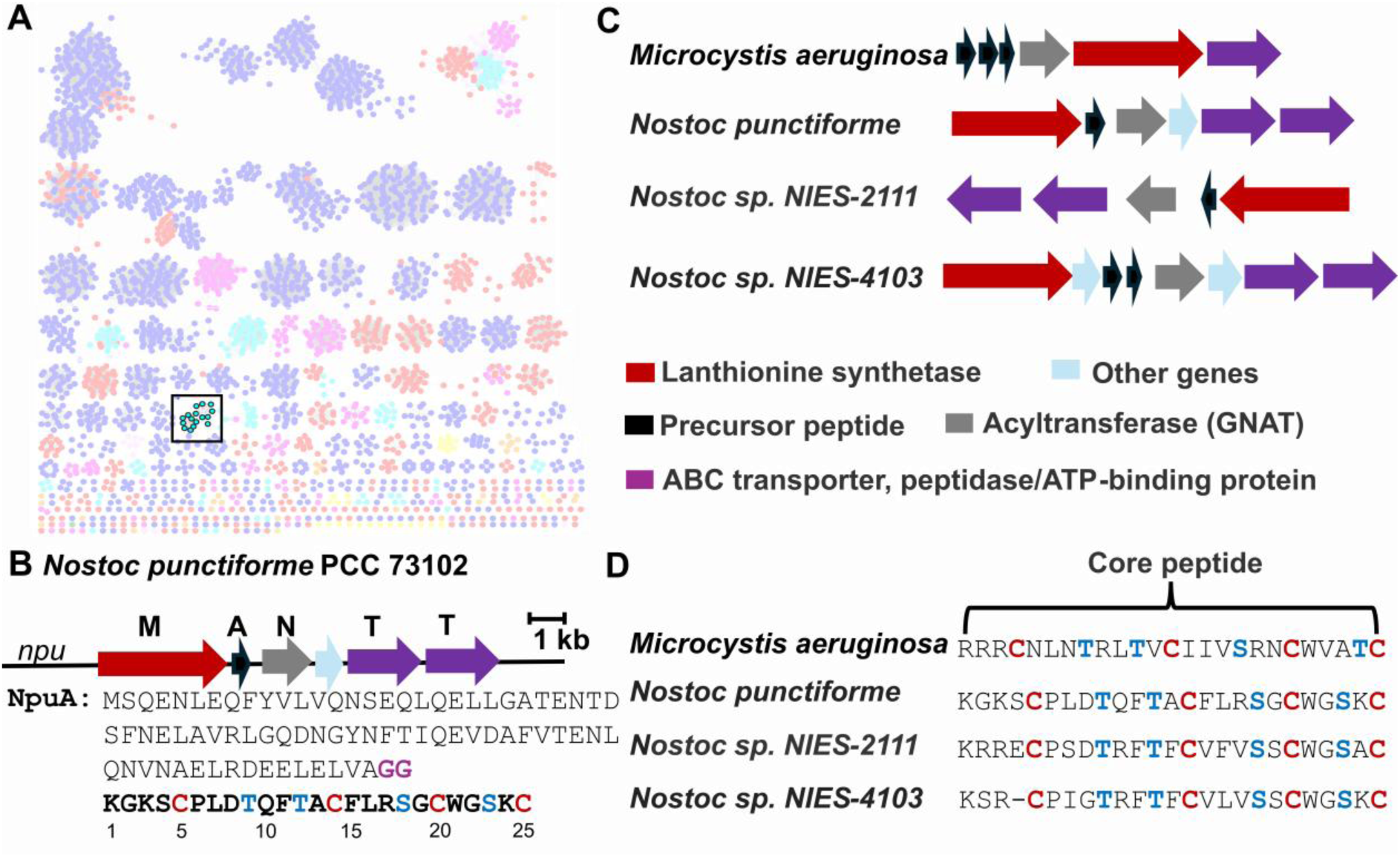
(**A**) Previously reported^39^ sequence similarity network (SSN) of predicted class II lanthipeptide precursor peptides generated with the Enzyme Function Initiative-Enzyme Similarity Tool.^40^ Each node represents a precursor sequence, and edges represent sequence similarity between peptides; Nodes are colored by phylum. The SSN was visualized with Cytoscape v3.10.^41^ Highlighted in cyan with black outline are the peptides that were previously designated cluster 65,^39^ which is the topic of this study. The cytoscape file is available as Supporting Information. (**B**) The nostolysamide BGC and the sequence of the precursor peptide NpuA. The predicted protease cleavage site for leader peptide removal is highlighted in purple and the core peptide is in bold. Residue numbering for the core peptide is indicated. (**C**) Representative BGCs associated with the 65 cluster in panel A, with genes colored according to predicted function. **(D**) Multiple sequence alignment of selected core peptides of the 65 cluster, highlighting conserved residues.

Co-expression of the bifunctional enzyme NpuM with N-terminally His6-tagged NpuA in *Escherichia coli* and subsequent purification of NpuA using immobilized metal affinity chromatography (IMAC) provided a fourfold dehydrated peptide. Subsequent *in vitro* removal of the leader peptide using LahT150,^44^ the protease domain of a homolog of NpuT, provided peptide **1** and demonstrated that the four dehydrations (−72.04 Da) occurred in the core peptide (**Fig. 2A, Fig. S1, S2**). To assess whether **1** was fully cyclized, an *N*-ethylmaleimide (NEM) assay was conducted to detect any free cysteines in the modified product;^46^ no NEM adducts were observed (**Fig. S3A**). Next, a dithiothreitol (DTT) assay was performed to test for the presence of dehydroamino acids in **1**. No adducts were observed (**Fig. S3A**). Collectively, these results indicate that **1** is fully cyclized.

**Figure 2.**
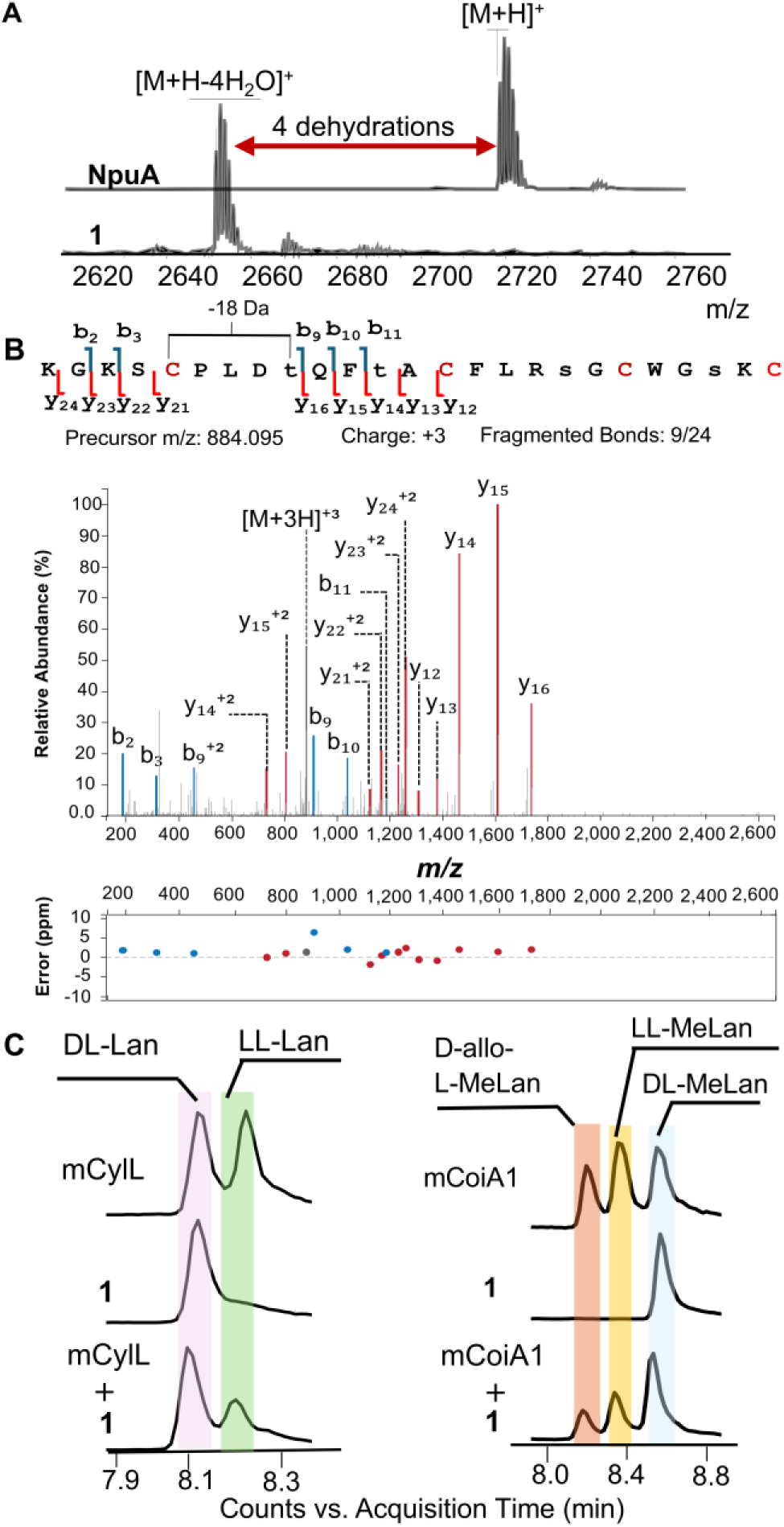
Structural characterization of NpuM-modified SUMO-NpuA. (**A**) MALDI-TOF mass spectra of NpuA before and after dehydration by NpuM, followed by cleavage with LahT150. NpuA, observed [M+H]^+^ m/z 2719.3; calculated m/z=2723.3. The formation of two disulfide bonds accounts for the observed 4 Da difference between calculated and observed ions. NpuA co-expressed with NpuM and cleaved with LahT150: observed m/z [M+H-4H O]^+^m/z=2651.2; calculated m/z=2651.2. (**B**) ESI-MS-MS data of **1** annotated with IPSA.^47^ The input sequence involved dehydrated residues at the positions indicated with lower-case letters. calculated m/z [M+3H-4H_2_O]^3+^ m/z=884.095, observed 884.095. (**C**) Marfey’s analysis of the Lan and MeLan rings in **1**. mCylL_L_ and mCoiA1 were used as standards, as reported previously.^48^

### Structural characterization of the modified peptides

As noted previously,^37^ production levels of modified peptide from co-expression of NpuA with NpuM in *E. coli* were low. We were able to somewhat improve production by using a small ubiquitin-like modifier (SUMO) fusion at the NpuA N-terminus. The product of co-expression of His_6_-SUMO-NpuA with NpuM again resulted in a fourfold dehydrated and fully cyclized product. While this improvement allowed structural characterization by tandem MS as discussed below, an initial screen of conditions for NMR analysis demonstrated insufficient signal dispersion and solubility of peptide **1** (e.g. **Fig. S4**). The peptide was therefore subjected to liquid chromatography-tandem mass spectrometry (LC-MS) analysis to determine the ring pattern through collision-activated dissociation (CAD) fragmentation. The fragmentation data was analyzed using the Interactive Peptide Spectral Annotator (IPSA),^47^ which generates predicted peptide fragments based on user-entered modifications. The tandem MS spectrum with IPSA annotations is presented in **Fig. 2B**. The fragmentation pattern is consistent with one MeLan ring formed from former Cys5 and Thr9. The very limited fragmentation between Cys14 and Cys25 made it difficult to determine the pattern of the other three rings and suggested overlapping rings (**Fig. 2B**). In addition, the fragmentation pattern and IPSA annotation suggested that Ser4 was not dehydrated.

To confirm this conclusion, we replaced Ser4 with Ala by site-directed mutagenesis. Heterologous expression of the SUMO-NpuA-S4A variant with NpuM yielded four dehydrations as observed for the wildtype (WT) peptide (**Fig. S5**), providing additional support that Ser4 is not dehydrated in WT NpuA.

To provide further insights into the apparent overlapping ring pattern involving Cys14, Cys20, and Cys25, an additional site-directed mutagenesis experiment was performed on the *npuA* gene to replace Cys25 with Ala. Heterologous co-expression of the SUMO-NpuA-C25A mutant with NpuM resulted in four dehydrations (**Fig. 3A**). NEM and DTT assays were carried out after proteolysis with LahT150 to ensure that the mutant peptide was fully cyclized (**Fig. S2B**). The DTT assay of NpuM-modified NpuA-C25A (**2**) showed a 154 Da adduct, indicating the presence of one dehydroamino acid, as expected, since one ring could not form as a result of the mutation. Furthermore, no NEM adduct was observed suggesting that the remaining Cys residues were all cyclized (**Fig. S3B**). Compound **2** was subjected to tandem MS analysis (**Fig. 3B**). Most observed ions were also present in the tandem MS spectrum of **1**, but two diagnostic new ions were observed with **2**. The y_12_^+2^ and y_13_^+2^ ions are most consistent with a methyllanthionine ring between Dhb12 and Cys25 in the WT compound that has been disrupted in the NpuA-C25A mutant. These two fragment ions together with a new y_6_ ion also suggest that Cys14 forms a ring with Ser18 and Cys20 with Ser23. As will be shown later, compound **2** was bioactive suggesting it was correctly cyclized.

**Figure 3.**
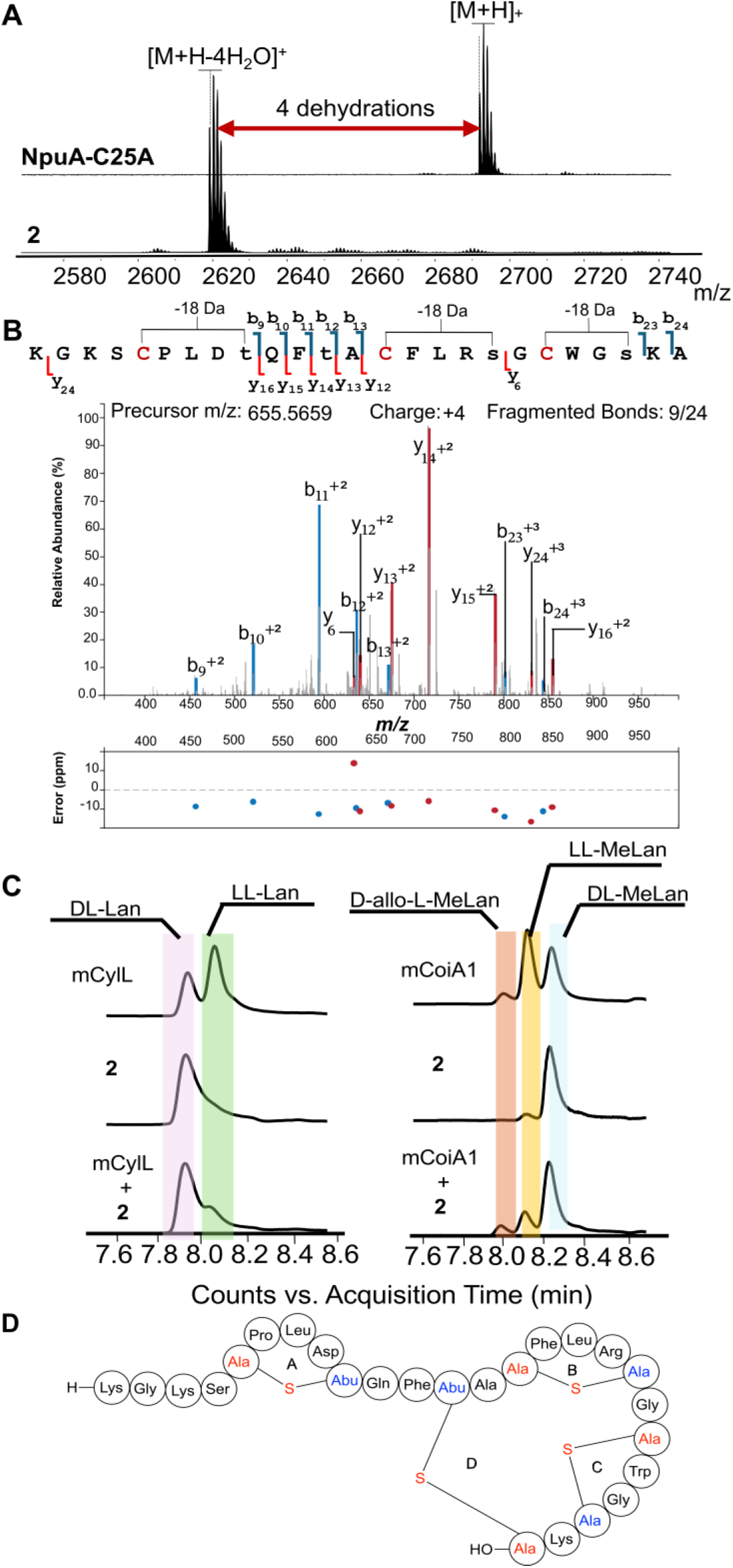
Structural characterization of NpuM-modified SUMO-NpuA-C25A. (**A**) MALDI-TOF mass spectra of NpuA-C25A before and after dehydration by NpuM, followed by cleavage with LahT150. NpuA-C25A, observed mass [M+H]^+^ m/z=2691.7; calculated m/z=2691.3. Product **2**, observed mass [M+H-4H_2_O]^+^ m/z=2619.4; calculated m/z=2619.2. (**B**) ESI MS-MS data of product **2**. Plot generated by IPSA showing b and y ions when the input sequence (lower-case letters) involved dehydrated residues at the indicated positions.^47^ Calculated m/z [M+4H-4H_2_O]^4+^ m/z=655.5659, observed 655.5712. (**C**) Marfey’s analysis of the Lan and MeLan rings in compound **2**. mCylL_L_ and mCoiA1 were used as standards.^48^ (**D**) Proposed ring pattern of nostolysamide C. The fragments of (methyl)lanthionines originating from Cys are colored red and the fragments originating from Ser/Thr are colored blue. Abu, 2-aminobutyric acid.

We also generated NpuA-C14A and NpuA-C20A variants and co-expressed them with NpuM in *E. coli*. Unlike NpuA-C25A, which was cleanly dehydrated four times, these variants generated a mixture of two, three and four dehydrations (**Fig. S6 and S7**). In addition, both peptides resulted in addition of glutathione (GSH) during co-expression and were incompletely cyclized as determined by NEM assays. These observations support our conclusion that Cys14 and Cys20 react with Dha18 and Dha23. GSH adducts to Dha residues during heterologous expression of lanthipeptides in *E. coli* has been observed previously,^38, 49–51^ including when cyclization is prevented either by mutagenic removal of the Cys residues that normally react with these Dha residues or by disruption of the cyclization enzyme.^49, 52^ Furthermore, current understanding of the post-translational modification process of lanthipeptides suggests that typically smaller rings are formed first,^53^ preorganizing the peptide for generation of larger rings.^54, 55^ When the formation of smaller rings is prevented by mutagenesis of substrate, both the dehydration and cyclization of larger rings is typically also negatively affected.^53^ Thus, for NpuA, disruption of formation of the B or C rings, likely also prevents formation of the D ring, resulting in GSH and NEM adducts.

### Stereochemistry of Lan and MeLan

The stereochemistry of the Lan and MeLan residues in peptides **1** and **2** was determined using advanced Marfey’s analysis. NpuM-modified SUMO-NpuA and SUMO-NpuA-C25A were subjected to hydrolysis under acidic conditions, followed by derivatization with the advanced Marfey reagent, Nα-(5-fluoro-2,4-dinitrophenyl)-L-leucinamide (L-FDLA).^48^ Extracted ion chromatograms (EICs) using the masses for derivatized Lan ([M –H]^-^ = 795.2373 Da) and MeLan [M–H]^-^ = 809.2530 Da) were monitored by LC-MS and compared to standards of DL– and LL-Lan and DL-, LL-, and *allo*-LL-MeLan derivatized in the same manner. Separate injections as well as co-injections demonstrated that the configuration of the Lan and MeLan residues in peptide **1** was DL (**Fig. 2C**).

The tandem MS data discussed above for the SUMO-NpuA-C25A mutant suggested the ring pattern shown in **Fig. 3B**. Previous NMR studies have validated that the ring patterns of native lanthipeptides determined from tandem MS analysis of mutants are often correct.^39, 56^ However, some studies have shown that use of fragmentation patterns for mutants can be misleading with respect to the ring pattern of the native sequence in cases of competing non-enzymatic cyclization, or non-native enzymatic cyclization when the normally highly ordered cyclization process is disrupted.^53, 57, 58^ Therefore, complementary data that provide support that mutants are correctly cyclized are important. Non-enzymatic (methyl)lanthionine formation often results in loss of stereochemical control.^59^ We therefore also determined the stereochemistry of the Lan and MeLan residues in NpuM-modified SUMO-NpuA-C25A (**2**). As shown in **Fig. 3C**, like peptide **1**, peptide **2** contains DL-Lan, and DL-MeLan, although a small amount of LL-MeLan was also observed. Together with the clean 4-fold dehydration, the stereochemical outcome suggests correct formation of the three (Me)Lan rings in mutant product **2**. Below, we will also show that this product has antifungal activity like product **1**, further supporting a native ring pattern. Therefore, our collective data support the ring pattern shown in Figure 3D for peptide **1**.

### Peptide 1 Exhibits Antimicrobial and Antifungal Activity

Peptide **1** was evaluated for antimicrobial activity against several Gram-positive, Gram-negative, and fungal strains using an agar diffusion assay. Compound **1** was dissolved in 5% aqueous dimethyl sulfoxide (DMSO) and spotted on agar plates. As a negative control, 5% aqueous DMSO was used, and kanamycin served as a positive control. Compound **1** demonstrated selective activity against several strains, as shown in **Fig. 4A**.

**Figure 4.**
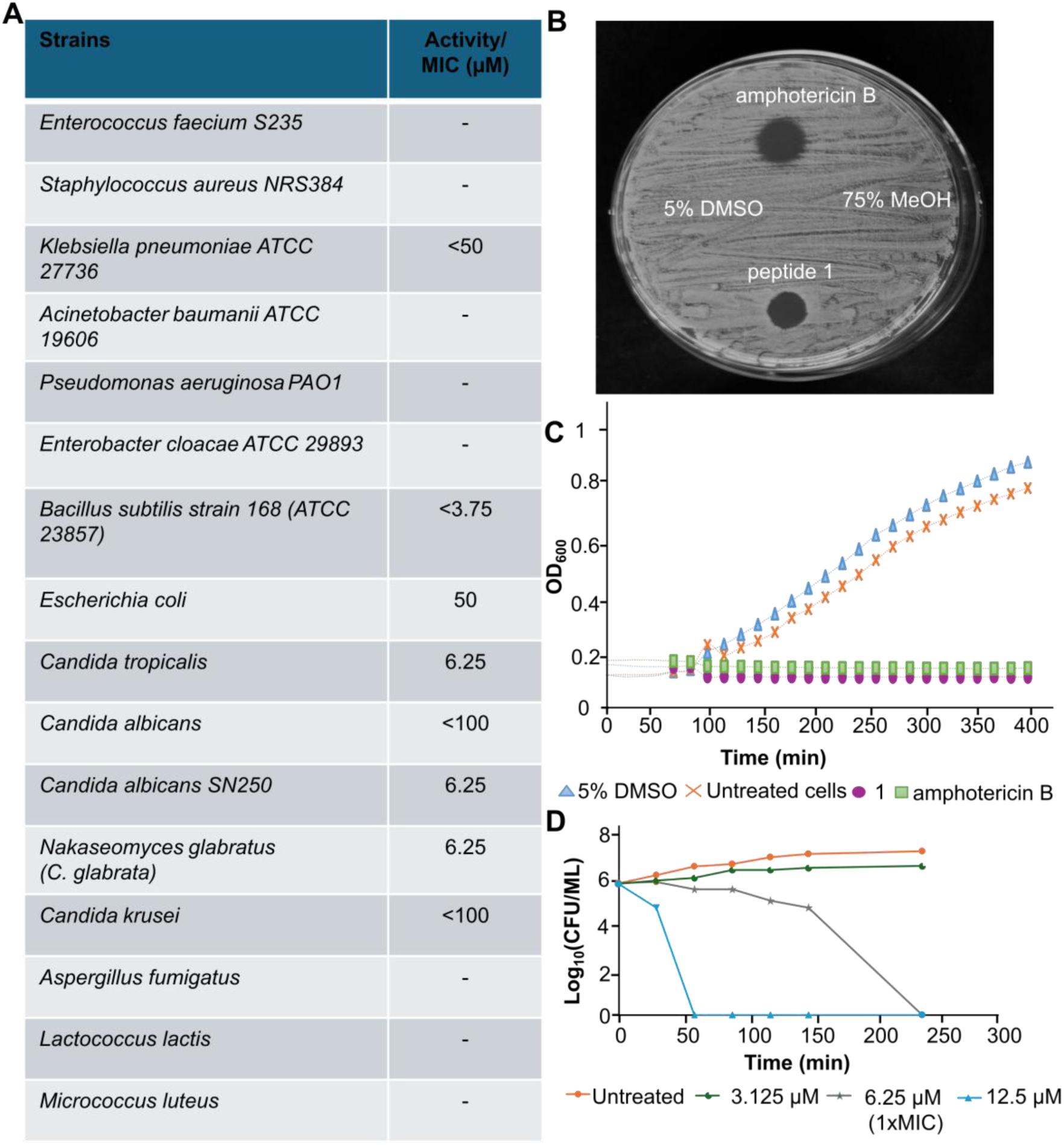
Antimicrobial activity of nostolysamide (1). (**A**) Activity of peptide **1** against various bacteria and fungal strains; – indicates no growth inhibition. (**B**) Agar diffusion assay with *Candida tropicalis* showing a clear zone of inhibition for peptide **1** (2 μL, 400 μM) and for the positive control amphotericin B (2 μL, 1 mM). No growth inhibition was observed for the negative controls 75% MeOH (2 μL) and 5% DMSO (2 μL). (**C**) Growth assay of *Candida tropicalis* displaying the optical density at 600 nm (OD₆₀₀) for cells that were treated with peptide **1** (100 μM), amphotericin B (200 μM), and negative controls. (**D**) Time-kill assay starting with 1 x 10^5^ colony-forming units (CFU) per mL of *Candida tropicalis* and monitoring the CFUs over 4 h using different concentrations of peptide **1**.

Peptide **1** inhibited the growth of *Bacillus subtilis* 168 (**Fig. S8**) and several *Candida* species (*Candida albicans* SN250*, Candida glabrata, Candida tropicalis*) at concentrations as low as 6.25 µM (**Fig. 4A, B; Fig. S9**). Using a broth dilution assay, weak activity was observed against some Gram-negative bacteria, including *E. coli* and *Klebsiella pneumoniae* ATCC 27736 (**Fig. S10**).

As noted in the introduction and discussed below, Piel and coworkers denoted the products of the *npu* BGC as nostolysamides, with acylated side chains of a Lys residue in these compounds; two molecules containing different acyl groups were termed nostolysamides A and B.^37^ Although peptide **1** does not feature an acyl group conjugated via an amide linkage to a Lys, we still consider it a congener of the products of the *npu* BGC and therefore will refer to it henceforth as nostolysamide C.

To assess the significance of the cyclic structure for activity, mutant **2** was also tested with both agar diffusion (**Fig. S11**) and broth dilution assays. Although the MIC value against *Candida tropicalis* increased more than 10-fold, the dehydrated and cyclized mutant peptide was still active. Given the rarity of antifungal activity, we interpret these data as showing that the mutant peptide is still correctly cyclized but that the ring involving Cys25 is important for antifungal activity. Nostolysamide C is the first known class II lanthipeptide to exhibit activity against *Candida* species, a common cause of fungal infection in humans. To probe whether the compound is fungistatic or fungicidal, *Candida tropicalis* was monitored over time after treatment, showing a decrease in colony forming units (CFUs) for both amphotericin B, as the positive control, and nostolysamide C compared to untreated cells (**Fig. 4D**), indicating that both molecules are fungicidal.

Class II lanthipeptides with antibacterial activity (lantibiotics) have diverse modes of action.^5^ Nisin and mersacidin-like compounds bind to the cell wall precursor lipid II in bacteria.^60–65^ Nostolysamide C was tested using a lipid II cycle-interfering assay that features a response regulator and sensor (LiaRS) assay. This system employs an engineered *Bacillus* strain in which a promoter controls lacZ gene expression, encoding the enzyme β-galactosidase, in response to interference with the lipid II cycle.^66^ When X-gal (a chromogenic substrate) was added to the medium, the positive control bacitracin produced the expected blue zone,^66^ but nostolysamide C and the negative control ampicillin did not (**Fig. S12**). These findings suggest that nostolysamide C does not bind to lipid II, which is consistent with its observed antifungal effect since fungi do not contain lipid II.

Given the antibacterial and antifungal activity displayed by nostolysamide C, the most likely target is the cell membrane. We therefore investigated whether nostolysamide C caused membrane depolarization by using a lipophilic cationic dye 3,3’-dipropylthiadixarbocyanine iodide (DiSC3(5)). In this assay, when the membrane of sensitive bacteria or fungi is depolarized, the fluorescence intensity of DiSC3(5) increases.^67, 68^ Addition of nostolysamide C indeed increased the fluorescence intensity like the positive control gramicidin^69, 70^ (**Fig. S13**), suggesting depolarization of the *Candida tropicalis* membrane.

### Characterization of the GNAT-acetyltransferase product

Nostolysamide C is the product of NpuM modification of NpuA followed by leader peptide removal. The observed antifungal and antimicrobial activity was somewhat surprising because an additional gene, *npuN*, was previously shown to add long chain fatty acids to Lys1 of NpuA.^37^ Hence, we had anticipated that the lipidation would be required for bioactivity. NpuN is a member of the GNAT family.^71^ Using NpuN as a query, we generated an SSN of related GNATs (**Fig. S14**). The BGCs associated with cluster 1 of this SSN contain putative RiPP precursor peptides containing either a nitrile hydratase like leader peptide (NHLP) or a Nif11-related leader peptide,^42, 43^ but they do not always encode a lanthionine synthetase (**Fig. S15**). A subset of these was shown in **Fig. 1C**, and a more extensive list is provided in **Fig. S15**. Most of these BGCs also encode an ABC transporter, suggesting that the acylated products are likely transported outside the cells and have extracellular functions.

Heterologous co-expression of *npuA* and *npuN* in *E. coli* resulted in peptides containing adducts of 237.3 Da and 252.3 Da (**Fig. 5A**); these same adducts were also observed for the dehydrated and cyclized peptide upon co-expression of all three genes—*npuA*, *npuM*, and *npuN* (**Fig. 5B**). The two fatty acid adducts, corresponding to hexadecenoyl and hydroxyhexadecenoyl groups, were also observed by Piel and coworkers who termed them nostolysamide A and B.^37^ Because of poor solubility, we were unable to purify the acylated products. We anticipated, based on previous data, that acylation occurred on Lys1.^37^ To confirm the regioselectivity of acylation and test how strict this selectivity is, site-directed mutagenesis was used to replace Lys1 and Lys3 in the core peptide of NpuA with alanine. For completeness, a double mutant NpuA-K1A/K3A was also constructed. When these mutant peptides were co-expressed with *npuN*, we observed an adduct only for NpuA-K3A (**Fig. S16**), demonstrating that acylation occurs on Lys1 and that Lys3 and Lys24 cannot substitute when Lys1 is absent.

**Figure 5.**
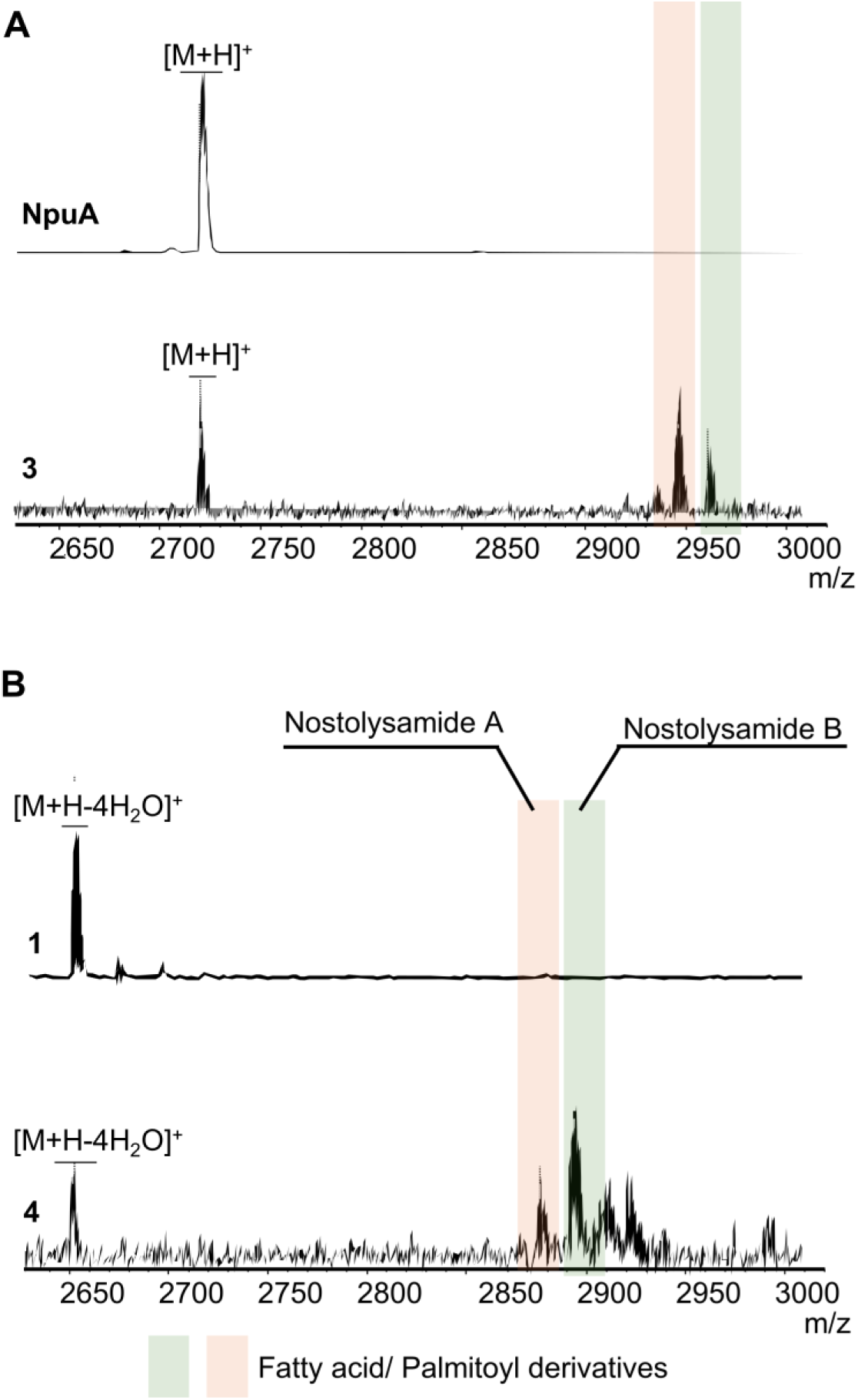
MALDI-TOF MS analysis of NpuN-modified peptide products. (**A**) MALDI-TOF mass spectra of NpuA and the peptide generated by coexpression of NpuA and NpuN, both treated with LahT150 *in vitro*; observed mass highlighted in orange and green, respectively: m/z 2957.2 and 2972.2. (**B**) MALDI-TOF mass spectra of the products of coexpression of NpuA and NpuM, and coexpression of NpuA, NpuM, and NpuN showing lipidation of the products; observed mass highlighted in orange and green m/z 2887.8 and 2903.7.

### NpuN displays substrate tolerance

The observed activity of NpuN was further evaluated *in vitro* by attaching a SUMO tag to its N-terminus to enhance solubility. His-SUMO-NpuN was expressed in *E. coli*, purified, and reacted with commercially available fatty acyl-CoAs and nostolysamide C (**1**) *in vitro* (**Fig. 6**). NpuN accepted acetyl-CoA, lauroyl-CoA, myristoyl-CoA, and oleoyl-CoA as substrates, resulting in the corresponding acylated nostolysamides (**Fig. 6A-E**). The peptide did not undergo complete modification and a mixture of acylated and non-acylated products was observed by MS that could not be purified because of the highly hydrophobic nature of the products, as also reported for the previously reported lipidated lanthipeptides kamptornamide and phaeornamide.^37^ The reaction was enzyme catalyzed because no acylation was observed when NpuN was withheld from these assays. Although the poor solubility prevented kinetic characterization, palmitoyl-CoA resulted in the highest conversion, consistent with the C16 products observed during the co-expression experiments discussed above and reported previously.^37^ Therefore, although it is possible that these acylated peptides may not represent the native products of *Nostoc punctiforme* PCC 73102, the observation of C16 products by co-expression and the apparent *in vitro* preference for palmitoyl-CoA suggest these are the physiological products. This hypothesis is also supported by the previous demonstration that the lipidated lanthipeptide phaeornamide produced in *E. coli* carried the same fatty acid as that detected in the native organism (*Pseudophaeobacter arcticus*).^37^

**Figure 6.**
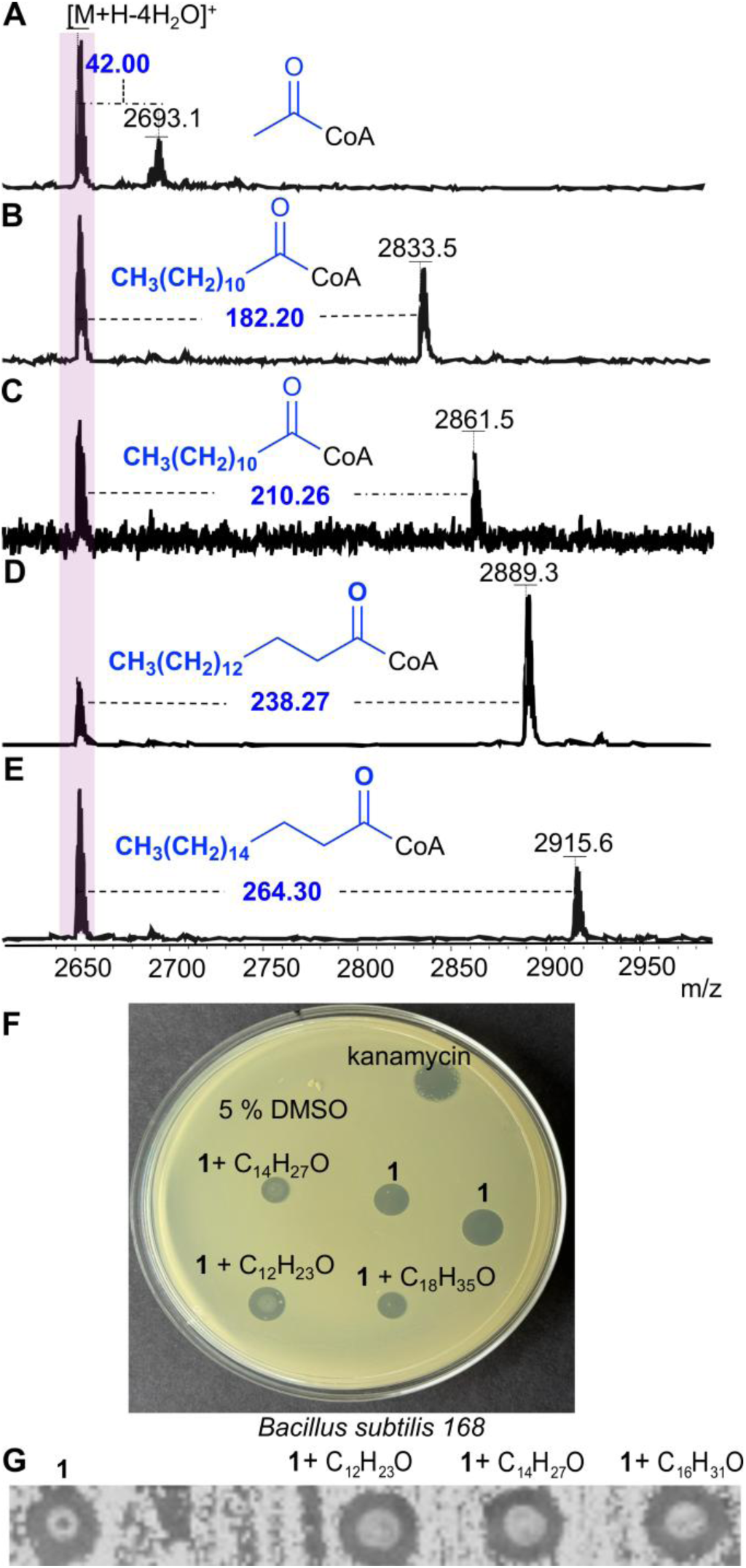
*In vitro* acylation of nostolysamide C by NpuN. (**A-E**) MALDI-TOF MS analysis of reactions containing acyl-CoA, His-SUMO-NpuN, and nostolysamide C. (**F**) Agar diffusion assay of palmitoylated, myristoylated, and laurylated nostolysamide against *B. subtilis* 168. (**G**) Antifungal activity of nostolysamides with or without acylation against *C. tropicalis*. GNAT enzymes involved in natural product acylation can serve self-resistance purposes or can enhance the biological activity of the compound.^36, 72, 73^ The *in vitro* acylated nostolysamides mixtures were therefore used for agar diffusion assays against *B. subtilis* and *C. tropicalis* to determine whether the activity would be enhanced or abolished. For three different acyl groups, antibacterial and antifungal activity was not significantly changed (Fig. 6F and 6G). The products were also subjected to the LiaRS assay, but no blue ring was observed (**Fig. S17B**). Hence, the function of the acylation of Lys1 is at present not clear.

### Predicted structure of NpuN

An AlphaFold3 model^74^ was generated for NpuN that revealed a tandem GNAT fold similar to other tandem GNAT proteins, a small group within the GNAT family (**Fig. S18A**).^75^ Compared to the typical single-domain GNATs,^26, 71, 76^ tandem GNAT proteins contain two GNAT-like domains within a single polypeptide. A DALI^77^ search of the AlphaFold3 model of NpuN retrieved the single domain tabtoxin resistance protein (TTR) complexed with acetyl coenzyme A as the closest structural homolog (PDB ID 1GHE)^78^ (**Fig. S18B**). Despite similar 3D folds, the sequence identity between NpuN and TTR is only 18.8%. (**Fig. S19A**). The acetyl-CoA binding site of GNATs is generated from a typical conserved P-loop motif [Q/R-X-X-G-X-G/A] that binds the diphosphate in acetyl-CoA.^71, 79^ Alignment of the sequence of TTR with NpuN identified the P-loop of NpuN in its N-terminal domain (**Fig. S19A**).

Virulence-induced protein F (VipF) from *Legionella hackeliae*, another tandem GNAT protein, contains P-loop sequences in both the N-terminal and a C-terminal domains.^80^ Based on an alignment of VipF and NpuN, the conserved P-loop motif is clearly present in the N-terminal domain of NpuN, but it is either absent or divergent in the C-terminal region (**Fig. S19**). In several tandem GNAT proteins studied including VipF the N-terminal domain is mainly non-catalytic and plays structural or regulatory roles, while the C-terminal domain is catalytic.^75, 80, 81^ However, exceptions occur for which the N-terminal domain is thought to be catalytic.^82^ A structural alignment of VipF and the AlphaFold model of NpuN suggests the acyl-CoA binding site in the N-terminal GNAT domain is mostly buried in both proteins. In contrast, the C-terminal domain, which in NpuN lacks the clear consensus sequence for acetyl-CoA binding, has an open, solvent-accessible binding groove for both proteins (**Fig. S18B**). An AlphaFold3 model of the complex of NpuA with NpuN also predicts that the peptide would bind preferentially to the C-terminal domain. In the model, potential catalytic residues Asp182 and His265 are positioned within about 5 Å of the Lys1 ε-amino group in the NpuA substrate (**Fig. S18C**). However, Asp and Tyr residues that were suggested to serve as a catalytic base to deprotonate the substrate and a catalytic acid to protonate the sulfur of the acyl-CoA in VipF,^80^ respectively, are also conserved in NpuN (**Fig. S18** and **S19**). Thus, experimental and/or structural data are needed to provide more insights into where catalysis occurs and which residues may be involved.

## Discussion

As part of our genome mining efforts using the automated FAST-RiPPs platform,^38^ a BGC from *Nostoc punctiforme* PCC 73102 was refactored for expression in *E. coli*. Low production levels and difficulty with product isolation, as well as the appearance of a literature report on the very same BGC,^37^ led us to not include our data in our previous report. However, the rapidly increasing interest in lipidated RiPPs and lipopeptides in general,^28, 31, 35, 36, 83–85^ and the discovery that the product displays antifungal activity prompted attempts to improve production to facilitate structure elucidation. Use of a fusion protein of the NpuA peptide and SUMO improved production sufficiently to carry out further biochemical characterization although yields remained low. Marfey’s analysis demonstrated that all four (Me)Lan residues have the DL configuration, and tandem MS analysis on the product formed by NpuM from SUMO-NpuA-C25A suggests the ring pattern shown in **Fig. 3D**. Determination of ring pattern for lanthipeptides with overlapping rings using mutagenesis can be perilous, but the observed clean formation of a 4-fold dehydrated and cyclized product with NpuA-C25A, the observation of the same stereochemistry of the crosslinks, and the observation of retention of some of the antifungal activity collectively support the notion that the NpuA-C25A variant is converted to a ring pattern that is the same as WT with the exception of the ring involving Cys25. The ring pattern in **Fig. 3D** shows that the ring involving Cys25 is the largest macrocycle in the proposed structure. Previous studies have shown that large rings are usually formed last during lanthipeptide synthesis, when the smaller rings that are formed first bring two residues that are distant in the primary sequence closer together.^54, 86^ In turn, disruption of the cyclization process involving rings that are formed last typically does not affect the formation of the other rings, explaining why NpuA-C25A still formed a bioactive product. The decrease in antifungal activity of the resulting variant does indicate the importance of the ring between former Cys25 and Thr12 for bioactivity.

Only a few cyanobacterial natural products are known to have activity against fungal species.^87^ Scytophycin is a broad-spectrum antifungal compound derived from *Scytonema*,^88^ and nostocyclamide, derived from *Nostoc*, has a narrow spectrum activity that targets *Candida albicans*.^89^ Another known cyanobacterial antifungal compound is calothrix A and B, isolated from *Calothrix,* which showed both antifungal and anticancer properties.^90^ The preliminary mode of action studies suggest that nostolysamide C targets the membrane, and the greatly reduced activity of the C25A variant indicates that the tetracyclic structure is important for activity.

As previously noted, pinensins were the first identified antifungal lanthipeptides. They are class I lanthipeptides and were isolated from a Gram-negative bacterium, *Chitinophaga pinensis*.^23^ Pinensins feature two methyllanthionine rings that do not have any sequence or structural similarity with the rings in nostolysamides (**Fig. S20**). Like the nostolysamides reported here, the antifungal activity of the pinensins is in the low micromolar range by disrupting membrane integrity.^23^ The pinensins are made by class I lanthipeptide machinery, including a LanC cyclase. LanC cyclases usually catalyze cyclization featuring a Cys residue that reacts with a Dha/Dhb residue that is located N-terminal to the Cys,^8^ and this is also the case for the pinensins. The nostolysamides are the first class II lanthipeptides reported to have antifungal activity. Most class II LanM enzymes catalyze cyclization with the same C-to-N directionality as in the pinensins.^8^ However, the ProcM-clade of class II enzymes from cyanobacteria,^9^ which includes NpuM, has been shown to catalyze cyclization in both directions involving Cys residues that are located both N– and C-terminal to the Dha/Dhb residues with which they react.^42, 56, 58^ For the ProcM-clade, the substrate sequence rather than the enzyme controls the cyclization pattern.^53, 91, 92^ Enzymes of the ProcM-clade are distinguished by having three Cys residues as ligands to the Zn^2+^ in the active site,^55^ rather than the two Cys and one His ligand set in class I LanC^10^ and most class II LanM enzymes.^93^ NpuM has the three Cys ligand set (**Fig. S21**), and indeed three of the four rings in the nostolysamides are made from a Cys that is located N-terminal to the partner Dha/Dhb. But unlike other ProcM-like enzymes in cyanobacteria, NpuM has only one substrate peptide rather than a set of different substrates with highly varying sequences.^42, 94, 95^

Our observations did not provide any support for lipidation improving the bioactivity of nostolysamides, in contrast to other reports on lipidation of peptide natural products. For instance, chemically attaching lipid groups to the RiPP aspergimycin significantly enhanced cytotoxic potency.^96^ Lipidation is a widely used strategy, both naturally and non-naturally, to improve the stability, cellular delivery, and bioactivity of peptide-derived natural products and we refer to reviews that discuss fatty acylation of various classes of natural products.^28, 36, 97–100^ NpuN is also not likely to be a self-resistance enzyme, a role fulfilled by several other GNATs.^72^ It is possible that an increase in bioactivity is delicately dependent on the precise molecular nature of the acyl group or the organisms targeted. Tests of these hypotheses will require isolation of the nostolysamides from the producing organism and testing against organisms that co-inhabit the same environment as *Nostoc punctiforme* PCC 73102, which was isolated from *Macrozamia* root. Given its complex life cycle involving vegetative cells, heterocysts, hormogonia and akinets,^101^ the production and function of the nostolysamides will require further investigation.

## Experimental Details

### Peptide and Protein Expression

Expression of His-NpuA, His-SUMO-NpuA, His-SUMO-NpuA/NpuM, His-SUMO-NpuA/NpuM/NpuN, His-SUMO-NpuA/NpuN, and His-LahT150, along with the NpuA variants (Supplementary Table S1), was performed using the same expression and purification method. For each sample, an overnight culture was grown from a single colony of *E. coli* BL21 (Tuner) transformed with the plasmid. This overnight culture was then diluted 1:100 into 1 L of LB broth and incubated at approximately 37 °C with shaking at 220 rpm until the culture reached an optical density of about 0.6. Co-expression with NpuN was adjusted so that cultures were grown at 30 °C until the desired OD_600_ was reached. The cultures were then placed on ice for about 30 min, followed by the addition of 0.5 mM isopropyl β-D-1-thiogalactopyranoside (IPTG). Afterward, the cultures were incubated at 18 °C with shaking at 180 rpm overnight for 16-21 h. Subsequently, the cell pellets were harvested by centrifuging at 4500 x g for 15 min.

### Generation of His-SUMO-NpuA variants

Site-directed mutagenesis was carried out using overlapping primers with the desired mutations (Supplementary Table S2). The following sets of polymerase chain reaction (PCR) conditions were established using the modified Q5 Hot Start High-Fidelity DNA polymerase protocol. A reaction volume of 30 μL was prepared, containing 2× Q5 Hot Start Master Mix, 0.5 μM of each primer (both forward and reverse), 10-50 ng of DNA template, and deionized water (DI H_2_O), adjusted to a total volume of 50 μL. The initial denaturation was carried out for 40 s at 98 °C, followed by 26 cycles of denaturation (9 s, 95 °C), annealing (20 s, 56-72 °C), and elongation (4 min, 71.5 °C), with a final elongation step (72 °C) for 6 min. After amplification, 1 μL of Dpn1 enzyme was added to each 50 μL reaction, and the mixture was incubated at 37 °C for 45 min. Successful amplification was confirmed via gel electrophoresis. Next, 3 μL of amplified product was added to 50 μL of chemically competent NEB10Beta electrocompetent cells. The cells were electroporated using an electroporation instrument and recovered with 900 μL of NEB10Beta recovery media for 1 h, shaking at 37 °C. After recovery, cells were pelleted, and ∼500 μL of supernatant was removed. The pellet was resuspended in the remaining media, and 50-100 μL was plated onto ampicillin plates. Plates were then incubated at 37 °C, after which single colonies were isolated and inoculated into 10 mL of LB containing ampicillin and grown overnight at 37 °C. Plasmid extraction was performed using Qiagen miniprep according to the manufacturer’s protocol, and the plasmids were sent for sequencing at Plasmidsaurus to confirm successful mutagenesis.

### Gibson Assembly^102^ using gBlocks

Synthetic DNA fragments (gBlocks, Integrated DNA Technologies) were designed with 10–30 bp overlapping regions complementary to the linearized vector backbone. The vector was linearized by PCR amplification using primers (Supplementary Table 2) and purified with a gel extraction protocol. The linearized vector (usually 0.02–0.5 pmol per fragment) and gBlocks (Supplementary Table 3) were combined to a final volume of 1.5–3 µL with Gibson Assembly 2X Master Mix (New England Biolabs) according to the manufacturer’s instructions. The reactions were incubated at 50 °C for 60 min. Electroporation-competent NEB10B cells were transformed with 2–5 µL of the assembly mixture. The transformed cells were recovered in NEB10B recovery medium for 1 h at 37 °C with shaking and then plated on LB agar containing the appropriate antibiotic. Colonies were picked, and plasmid DNA was extracted using a miniprep protocol. The plasmid sequences were verified by Plasmidsaurus to confirm the correct assembly of the plasmids used in the study (Supplementary Table 1).

### Purification of peptides

Cell pellets were harvested and resuspended in Lysis buffer (20 mM Na_2_HPO_4_, 10 mM imidazole, 300 mM NaCl and 6 M guanidine hydrochloride, Gn-HCl, pH 7.5). This mixture was sonicated (40% amplitude, 4.4 s on, 5.5 off) for 6 min. The lysate was centrifuged at 10,000 x g for 30 min. The supernatant was loaded onto an IMAC column equilibrated with Lysis buffer, using 0.5 mL of Ni resin per gram of cell pellet. This step was followed by washing with 2–3 column volumes (CV) of Lysis buffer followed by Wash buffer 1 (20 mM Na_2_HPO_4_, 10 mM imidazole, 300 mM NaCl and 4 M Gn-HCl). The column was washed with 2-3 CV of Wash buffer 2 (20 mM Na_2_HPO_4_, 30 mM imidazole, 300 mM NaCl, pH 7.5) before eluting the peptide with 2-3 CV of Elution buffer (20 mM Na_2_HPO_4_, 1 M imidazole, 300 mM NaCl, pH 7.5).

### Purification of Protease His-LahT150, His-NpuN, and His-SUMO-NpuN

Cell pellets were resuspended in protein Start buffer (20 mM Tris-HCl, 10 mM imidazole, 300 mM NaCl and 10% glycerol, pH 8.0) containing 4 mg/mL of lysozyme. This mixture was incubated at 4 °C, then lysed via sonication (40% amplitude, 4.4 s on, 5.5 off) for 6 min. The lysate mixture was centrifuged at 10,000 x g for 30 min. The supernatant was loaded onto an IMAC column equilibrated with protein Start buffer, using 0.5 mL of Ni resin per gram of cell pellet. This step was followed by washing with 2–3 CV of protein Start buffer. The column was washed with 2-3 CV of protein Wash buffer (20 mM Tris-HCl, 30 mM imidazole, 300 mM NaCl and 10% glycerol, pH 8.0) before eluting the protein with 2-3 CV of protein Elution buffer (20 mM Tris-HCl, 300 mM imidazole, 300 mM NaCl and 10% glycerol, pH 8.0). Fractions were analyzed by sodium dodecylsulfate polyacrylamide gel electrophoresis using a 4–20% Tris gel at 150 V. The elution fraction was collected, concentrated, and the buffer exchanged to protein Storage buffer (20 mM Tris-HCl, 500 mM NaCl, and 10% glycerol, pH 8.0) using a 3 kDa MWCO Amicon filter. The fraction was then aliquoted and stored at −80 °C.

### *In vitro* cleavage with LahT150

LahT150 cleavage reactions were performed at 25 °C for 18 h on the modified precursor peptides to obtain the modified core peptides for further purification and analysis. The elution buffer from peptide purification was exchanged to 50 mM Tris-HCl, 100 mM NaCl (pH 8.0) using an Amicon tube with a 3 kDa cutoff. LahT150 (2.5 mg/mL, approximately 136 µM) was added at 100 µL/L to _the_ sample. The LahT150 assay was quenched by adding 0.1% v/v trifluoroacetic acid (TFA) in H_2_O, and the mixture was centrifuged to remove precipitates. The supernatant was used for further purification.

### HPLC purification of modified peptides

After IMAC purification, the modified and unmodified LahT150-cleaved peptides were subjected to further purification on an Agilent 1260 II Infinity HPLC. The peptides were purified by RP-HPLC using a Hypersil GOLD C8 column (5 μm, 175 Å, 250 mm × 4.6 mm) with a linear gradient from 2% solvent B (100% MeCN with 0.1% TFA) in solvent A (0.1% TFA in water) to 100% solvent B over 33 min. The absorbance was monitored at 220 and 280 nm. Fractions containing the cleaved, modified core were identified by MALDI-TOF MS and lyophilized for further characterization.

### NEM Assay

A literature procedure^46^ was adapted. A 50 μL reaction was set up with 25-50 μM peptide dissolved in 50 mM Tris-HCl, pH 7.5 (25 μL), and 0.3 mM of fresh 1.0 mM stock solution of tris(2-carboxyethyl)phosphine (TCEP) (20 μL) that was incubated for 30 min at room temperature before 500 mM NEM (5 μL) was added. The reaction was incubated at room temperature for 3 h before using a C18 ziptip to prepare the sample for analysis by MALDI-TOF MS. For every free Cys in the peptide, a mass shift of +125 Da is expected.

### DTT Assay

A literature procedure^46^ was adapted. The peptide was dissolved in 50 μL of 100 mM Tris-HCl buffer, pH 7.5, to obtain a final concentration of 100 μM. *N*, *N*-diisopropylethylamine (DIPEA) and DTT were added to give final concentrations of 400 mM and 500 mM, respectively. The 50 μL reaction was incubated at room temperature for 3 h. After desalting with a C18 ziptip, the reaction was analyzed by MALDI-TOF MS.

### High-Resolution Tandem Mass Spectrometry

The HPLC-purified peptides were injected onto a ThermoFisher Scientific Orbitrap Fusion electrospray ionization (HR-ESI) mass spectrometer using an Advion TriVersa Nanomate 100. The parameters used for MS(/MS) data acquisition included a resolution of 100,000 in positive mode, an isolation width of 0.5–2 m/z (MS/MS), and a normalized collision energy of 20, 25, and 30 (MS/MS). Data analysis was performed using the Xcalibur software.

### Marfey’s Analysis

For Marfey analysis, the two standards mCylL_L_ and mCoiA1, along with NpuA and mutants, were purified after heterologous expression in *E. coli* as described previously,^48^ and derivatized with slight modifications. Dried peptides (∼100 μg) were dissolved in 1 mL of 6 M deuterium chloride (DCl) in D_2_O for hydrolysis using a long glass vial. N_2_ gas was bubbled in the glass vials for 1 min. The reaction was heated to 120 °C and stirred for 20 h. Each reaction product was then dried using a rotary evaporator before being placed on a lyophilizer for 1 h to remove any remaining DCl. This step was followed by adding 0.6 mL of 0.8 M NaHCO_3_ in H_2_O and 0.4 mL of 4 mg/mL of Nα-(5-fluoro-2,4-dinitrophenyl)-l-leucinamide (L-FDLA) in acetonitrile. Next, the glass vials were stirred in the dark at 67 °C for 3 h. Subsequently, 100 μL of 6 M HCl was added to the derivatization product, and the mixture was vortexed. The mixture was placed on a lyophilizer to dry, and the residue was resuspended in 500 μL of acetonitrile. The resuspension mixture was centrifuged at high speed for 10 min, and the supernatant was collected and analyzed on an Agilent 6545 LC/Q-TOF instrument using a Kinetex 1.7 μm F5 100 Å, LC column (100 × 2.1 mm; Phenomenex; part no.: 00D-4722-AN) at a constant flow rate of 0.4 mL/min. The eluting system consisted of a gradient of 5-80 % of solvent B (MeCN with 0.1% formic acid) in solvent A (water with 0.1% formic acid). Chromatographic separation was obtained for each derivatized diastereomer and the identity of the NpuA samples was confirmed by co-injection and extracting the exact mass for derivatized MeLan (m/z 809.25299) and Lan (m/z 795.23734).

### Agar Well Diffusion Assays to Determine Antimicrobial Activity

Agar well diffusion growth-inhibition assays were conducted following a biosynthesis protocol from our previous report.^38^ Peptides were dissolved in 5% DMSO to achieve a starting concentration of 0.5 mg/mL. Agar plates were prepared by making 1% agar with the desired media, approximately 20 mL for the bottom agar. The top agar was prepared by adding 200 μL of the overnight cell culture to 20 mL of melted agar cooled to about 55 °C. The cultured agar was then poured onto the bottom agar to solidify. The dissolved peptide was directly spotted onto the solidified plate. For the fungi agar assay, sterile cotton swabs or Q-tips were used to streak the overnight fungi culture on the plate before spotting the dissolved peptide. The dried plates were incubated at 37 °C for 18 h, and the presence or absence of zones of growth inhibition determined the activity. The negative control was 5% DMSO, and the positive control was kanamycin for the bacterial assay or amphotericin B for the fungal assay.

### Minimal Inhibitory Concentration (MIC) Determination

The method was modified from a previously described procedure.^103^ The MIC was determined using the standard doubling dilution method with Lysogeny broth (LB), Mueller-Hinton broth (MHB), or Roswell Park Memorial Institute (RPMI) 1640 for fungal species. All tests with the examined wells were performed with three biological replicates. Overnight cultures were inoculated and grown for an additional 4 h before adjusting the bacterial culture to approximately 10^5^ CFU per mL. The assay was set up with wells containing various concentrations, ranging from 0.78 μM to 200 μM of the tested peptide. A positive control was included with kanamycin or amphotericin B, a negative control with just 5% DMSO, and a row with only medium. For the antifungal MIC against *Candida* species, as previously described,^103^ an inoculum of 1 × 10^5^ cells in 190 μL of RPMI 1640 medium was added to 96-well polystyrene microplates with different concentrations of the tested peptide and amphotericin B. After incubation at 37 °C for 18 h, the OD_600_ was measured using a plate reader. The MIC was defined as the lowest concentration at which the tested peptide caused growth inhibition.

### LiaRS assay

The method followed the previously described protocol with modifications.^66^ An overnight culture of *B. subtilis* BSF2470 was grown to an OD_600_ of 1.0 before dilution with 0.7% molten LB agar containing 20 mg/mL X-Gal in dimethylformamide. The molten agar was overlayed on a solidified bottom agar. The plates were allowed to cool and dry on the bench at room temperature for 30 min. Then samples were spotted directly on the plates in the following manner: 100 mg/mL ampicillin (2 µL – negative control), 35 mM bacitracin (2 µL – positive control), 2 µL of 5% DMSO, and 2 µL of 200 µM peptide **1**. Plates were left for 45 min before being placed in a 37 °C incubator for 18 h. A blue ring around the zone of inhibition indicates the compound targets the bacterial cell wall or membrane, including through lipid II associated processes.

### Time killing assay of peptide 1 against *Candida tropicalis*

Time-killing assays were performed according to a previously described protocol, with modifications.^104^ Briefly, a growth curve was generated in the absence of the tested peptide to determine the doubling time of *Candida tropicalis*. The experiment was conducted in triplicate. An overnight culture was inoculated and diluted 100-fold in RPMI 1640 medium. The cells were incubated at 37 °C, then grown for an additional 4 h before being incubated at 37 °C to an OD_600_ of 1.0. The cell concentration was adjusted to 5 × 10^5^ CFU/mL. Fungi were then incubated at half the MIC and at the MIC of peptide **1**. Cells treated with 5% DMSO were used as a control. At the 4-hour time point, 10 μL was taken from each sample, diluted 1000-fold in RPMI 1640, and plated on Sabouraud Dextrose agar plates. After overnight incubation at 37 °C, colonies on the plates were counted and expressed as colony-forming units per mL to generate the plot.

### DiSC3(5) Assay^67^

Cells were grown in potato dextrose (PD) medium until logarithmic growth was obtained. Cells were diluted to an OD600 of 0.2 in pre-warmed PD supplemented with 0.5 mg/mL BSA. Fluorescence measurements were performed using a Tecan Plate Reader Infinite F200 with an excitation wavelength at 622 nm and emission at 670 nm. Membrane depolarization was indicated by an increase in fluorescence intensity due to dye release into the extracellular medium. Cells (135 mL) were transferred to a microtiter plate and the fluorescence was recorded for 2–3 min in order to obtain values for medium and cell background. After obtaining a baseline, DiSC3(5) dissolved in DMSO (100 μM) was added to each well to a final concentration 1 μM DiSC3(5) and 1% DMSO. Maintaining 1% DMSO was critical for good solubility and fluorescence of the dye. Fluorescence was measured until a stable signal intensity was achieved, followed by addition of the compound of interest. To separate wells for the positive control 1 μM of channel-forming peptide gramicidin S was used. Nostolysamide C was used for the tested compound; 10 μL of 50 μM and 100 μM solutions of **1** were added to separate wells, and 5% DMSO was used as a negative control. Fluorescence readings were recorded over 60 min.

### *In vitro* reaction with His-SUMO-NpuN

The NpuA peptide was dissolved in 5% DMSO to a concentration of 50 μM. His-SUMO-NpuN was added to a final concentration of 17 μM, along with 200 μM of fatty acyl-CoA. Tris-HCl at pH 8.5 was added to make the total reaction volume 100 μL. The reaction was incubated at 37 °C and analyzed by MALDI-TOF MS after 16-18 h.

## Supporting Information

Figures S1-S21 showing mass spectrometry and bioactivity data, and AlphaFold models, and Tables S1-S5 with gene sequences and accession numbers, primers, and plasmids used (PDF).

## AUTHOR INFORMATION

### Corresponding Author

Wilfred A. van der Donk − Department of Chemistry and Howard Hughes Medical Institute, University of Illinois at Urbana-Champaign, Urbana, IL, United States; orcid.org/0000-0002-5467-7071.

### Authors

Enleyona Weir – Department of Chemistry and Howard Hughes Medical Institute, University of Illinois at Urbana-Champaign, Urbana, IL, United States.

Ivan Anterola – Department of Chemistry, University of Illinois at Urbana-Champaign, Urbana, IL, United States.

## Supporting information

Cytoscape files

## Acknowledgements

The authors thank Youran Luo for help with Marfey’s analysis, Dr. Chengyou Shi for the construction of the original plasmid encoding NpuA and NpuM, and Dr. Chandrashekhar Padhi for helpful discussions regarding fungal assays.

## Notes

The authors declare no competing financial interests.

## Data availability

Raw data are deposited on Mendeley and will be released upon publication. Weir, E.; van der Donk, W.A. (2026), Data associated with “Structure and biosynthesis of nostolysamides”, Mendeley Data, V1, doi: 10.17632/bczwrs7nr9.1.

---

| **Accession number** | **NCBI accession** | **Function** |
| --- | --- | --- |
| NpuA | WP_012412980.1 | Precursor peptide |
| NpuM | WP_012412979.1 | Lanthipeptide synthetase |
| NpuN | WP_012412981.1 | GNAT acyl transferase |

