## Supporting Information for "Structure, biosynthesis, and bioactivity of nostolysamides"

Table S1: Plasmids used in this study ........ S2
Table S2. Primers used in this study ........ S3-S4
Table S3. gBlocks used in this study ........ S4
Table S4. Accession number associated with the proteins used ........ S5
Table S5. Strains associated with cluster 65 ........ S5
Figure S1. LC-QTOF-ESI/MS profile of nostolysamide C ........ S6
Figure S2. Reversed-phase HPLC traces of purified nostolysamide C ........ S7
Figure S3. MALDI-TOF mass spectra of the NEM and DTT assay of **1** and **2** ........ S8
Figure S4: $^1$H NMR spectrum of nostolysamide C ........ S9
Figure S5. Characterization of SUMO-NpuA-S4A modified by NpuM ........ S10
Figure S6. MALDI-TOF MS of NpuA-C14A co-expressed with NpuM ........ S11
Figure S7. MALDI-TOF MS of NpuA-C20A co-expressed with NpuM ........ S12
Figure S8. Bioactivity assay with Compound **1** against gram-positive bacteria ........ S13
Figure S9. Bioactivity assay with nostolysamide C against Candida species ........ S13
Figure S10. Bioactivity assay with Compound **1** against gram–negative bacteria ........ S14
Figure S11. Bioactivity assay with Compound **2** against *Candida tropicalis* ........ S14
Figure S12. Compound **1** does not target lipid II ........ S15
Figure S13. Membrane depolarization measured by DiSC$_3$(5) assay ........ S16
Figure S14. Sequence Similarity Network (SSN) of GNAT enzymes ........ S17
Figure S15. A representative subset of genome neighborhoods from Cluster 1 ........ S18
Figure S16. MALDI-TOF MS of the lysine mutants ........ S19
Figure S17. Analysis of acylated product using SUMO-NpuN ........ S20-21
Figure S18. Structural comparison of GNAT protein ........ S22-23
Figure S19. Sequence alignments of the GNAT protein ........ S24-25
Figure S20. Structure of pinensins ........ S25
Figure S21. Sequence alignment of NpuM and ProcM ........ S25
References ........ S26

**Supplementary Table S1: Plasmids used in this study**

| **Plasmids** | | |
|---|---|---|
| pET-$His_6$-SUMO-NpuA_NpuM | pETduet derivative, expression $His_6$-SUMO-NpuA_NpuM, Ampicillin | This work |
| pRSFduet-$His_6$-NpuM | pRSFduet derivative, expression $His_6$-NpuM, kanamycin | This work |
| pRSFduet-$His_6$-NpuN | pRSFduet derivative, expression $His_6$-NpuN, kanamycin | This work |
| pET-$His_6$-SUMO-NpuA | pETduet derivative, expression $His_6$-SUMO-NpuA, Ampicillin | This work |
| pRSFduet-$His_6$-SUMO-NpuN | pRSFduet derivative, expression $His_6$-SUMO-NpuN, kanamycin | This work |
| pET-$His_6$-SUMO-NpuA_K1A | pETduet derivative, expression $His_6$-SUMO-NpuA_K1A, Ampicillin | This work |
| pET-$His_6$-SUMO-NpuA_K3A | pETduet derivative, expression $His_6$-SUMO-NpuA_ K3A, Ampicillin | This work |
| pET-$His_6$-SUMO-NpuA_K1A/K3A | pETduet derivative, expression $His_6$-SUMO-NpuA_ K1A/K3A, Ampicillin | This work |
| pET-$His_6$-SUMO-NpuA_C25A | pETduet derivative, expression $His_6$-SUMO-NpuA_ C25A, Ampicillin | This work |
| pET-$His_6$-SUMO-NpuA_S18A | pETduet derivative, expression $His_6$-SUMO-NpuA_ S18A, Ampicillin | This work |
| pET-$His_6$-SUMO-NpuA_S4A_NpuM | pETduet derivative, expression $His_6$-SUMO-NpuA_S4A_NpuM, Ampicillin | This work |

**Supplementary Table S2: Primers used in this study**

| Template | Primers | Nucleotide sequence (5` to 3`) | Plasmid produced |
|---|---|---|---|
| pET-28-His6-NpuA-NpuM | NpuA_65_SUMO_F | gatcggatcctATGTCCCAGGAGAACC | pET-His6-SUMO-NpuA |
| | NpuA_65_SUMO_R | gctcgaattcttaGCATTTGGAGCCCC | |
| pET- His6-SUMO | pET-v-SUMO-F | TCCAAATGCtaagaattcgagctcggc | |
| | pET-v-SUMO-R | ATaggatccgatccaccaatctgttctctgt | |
| pET- His6-SUMO-NpuA | pET-v-SUMO-NpuA-NpuM-F | tgagcggaattcgagctcggc | pET-His6-SUMO-NpuA_NpuM |
| | pET-v-SUMO-NpuA-NpuM-R | ctagaccgttaGCATTTGGAGCCCC | |
| pET-28-His6-NpuA-NpuM | pET-gM-SUMO-F | CAAATGCtaacggtctagcataacccettg | |
| | pET-gM-SUMO-R | ctcgaattccgctcaaaaaacccctca | |
| pET- His6-SUMO-NpuA | NpuAS4A-F | GAGGTAAAGGAAAGgctTGTCCGCTGGACACGC | pET-His6-SUMO-NpuA_S4A_NpuM |
| | NpuAS4A-R | GCGGACAagcCTTTCCTTTACCTCCCGCAAC | |
| pET- His6-SUMO-NpuA | NpuAC25A-F | GGGGCTCCAAAgcttaagaattcgagctcggcgc | pET-His6-SUMO-NpuA_C25A |
| | NpuAC25A-F | agctcgaattcttaagcTTTGGAGCCCCAGCAACCAGAG | |
| pET- His6-SUMO-NpuA | NpuAS18A-F | CCTGCGCgctGGTTGCTGGGGCTC | pET-His6-SUMO-NpuA_S18A |
| | NpuAS18A-R | CCCAGCAACCagcGCGCAGGAAGCAAG | |
| NpuN-gblock | NpuN-F | GCAGGCGCGCCGAGC | pRSFduet-His6-NpuN |
| | NpuN-R | CCCTGTAGAAATAATTTTGTTTAACTTTAATAAGGAGATATACCATGGGC | |
| pRSF-His6 | pRSF-v-NpuN-F | GGCGCGCCTGCAGGTC | |
| | pRSF-v-NpuN-R | CTCCTTATTAAAGTTAAACAAAATTATTTCTACAGGGGAATTGTTATCCG | |
| pRSF-His6 | pRSF-v-NpuM-F | CTGTGGGAAtaaAGCCAGGATCCGAATTC | pRSFduet-His6-NpuM |
| | pRSF-v-NpuM-R | GCGTTCTTCCATGTGGTGATGATGGTGA | |

| | | | |
|---|---|---|---|
| pET-28-$His_6$-NpuA_NpuM | NpuM-F | CATCACCACATGGAAGAACGCGAAATGATCAAT | |
| | NpuM-R | GGATCCTGGCTttaTTCCCACAGAAGCAC | |
| pRSF_SUMO-TEV $His_6$ | pRSF-v-SUMO_NpuN-F | ggctaatgcggccgcataatgct | pRSFduet-$His_6$-SUMO-NpuN |
| | pRSF-v-SUMO_NpuN-R | gttcagggtatacattccctggaagtacaggttttc | |
| pRSFduet-$His_6$-NpuN | NpuN-F | cctgtacttccagggaatgtataccctgaacatt | |
| | NpuN-R | gttcagggtatacattccctggaagtacaggttttc | |

**Supplementary Table S3: gBlocks used in this study**

| | |
|---|---|
| NpuA_Tev_K1A | ggatcggatcctGAAAATTTATATTTTCAGAGCATGTCCCAGGAGAACCTGGAACAGTTTTACGTCCTCGTGCAAAACTCAGAACAATTACAAGAGCTGTTGGGCGCTACGGAAAACACAGATTCATTTAATGAACTCGCAGTGCGTTTAGGCCAGGATAATGGGTATAACTTTACCATCCAAGAAGTTGATGCTTTCGTCACCGAAAATCTGCAGAACGTTAACGCGGAACTGCGTGATGAAGAACTGGAACTGGTTGCGGGAGGTgcgGGAAAGTCGTGTCCGCTGGACACGCAATTCACGGCTTGCTTCCTGCGCTCTGGTTGCTGGGGCTCCAAATGCtaagaattcgagct |
| ENW_Tev_K1A_K3A | ggatcggatcctGAAAATTTATATTTTCAGAGCATGTCCCAGGAGAACCTGGAACAGTTTTACGTCCTCGTGCAAAACTCAGAACAATTACAAGAGCTGTTGGGCGCTACGGAAAACACAGATTCATTTAATGAACTCGCAGTGCGTTTAGGCCAGGATAATGGGTATAACTTTACCATCCAAGAAGTTGATGCTTTCGTCACCGAAAATCTGCAGAACGTTAACGCGGAACTGCGTGATGAAGAACTGGAACTGGTTGCGGGAGGTgctGGAGCTTCGTGTCCGCTGGACACGCAATTCACGGCTTGCTTCCTGCGCTCTGGTTGCTGGGGCTCCAAATGCtaagaattcgagct |
| ENW_Tev_K3A | ggatcggatcctGAAAATTTATATTTTCAGAGCATGTCCCAGGAGAACCTGGAACAGTTTTACGTCCTCGTGCAAAACTCAGAACAATTACAAGAGCTGTTGGGCGCTACGGAAAACACAGATTCATTTAATGAACTCGCAGTGCGTTTAGGCCAGGATAATGGGTATAACTTTACCATCCAAGAAGTTGATGCTTTCGTCACCGAAAATCTGCAGAACGTTAACGCGGAACTGCGTGATGAAGAACTGGAACTGGTTGCGGGAGGTaaaGGAGCTTCGTGTCCGCTGGACACGCAATTCACGGCTTGCTTCCTGCGCTCTGGTTGCTGGGGCTCCAAATGCtaagaattcgagct |

**Supplementary Table S4:** Accession numbers associated with proteins used in this study

| Accession number | Protein |
|---|---|
| NpuA | WP_012412980.1 |
| NpuM | WP_012412979.1 |
| NpuN | WP_012412981.1 |

**Supplementary Table S5:** Strains associated with cluster 65 in the SSN (Fig. 1, main text) and their precursor peptides. Conserved residues that are predicted to be modified by class II lanthipeptide synthetases are highlighted.

| Strain | Precursor sequence |
|---|---|
| *Nostoc sp. NIES-2111* | MNNSNLQNFYTLIQNSQELQAQLGAADSPESFAETAVRLGE ENGYSFTTEDVNTFISQQRSRANAELSEADLEAVAGGKRRE CPSDTRFTFCVFVSSCWGSAC |
| *Nostoc punctiforme* | MSQENLEQFYVLVQNSEQLQELLGATENTDSFNELAVRLGQ DNGYNFTIQEVDAFVTENLQNVNAELRDEELELVAGGKGKS CPLDTQFTACFLRSGCWGSKC |
| *Microcystis aeruginosa* | MSQPDVERFYEIARNSEELQAMLGAAEDRDSFVETAVRLG QENGCNFTLDEANEFLNTKKGDAELSEQELEAVAGGRRRC NLNTRLTVCIIVSRNCWVATC |
| *Nostoc sp. NIES-4103* | MSKENLEQFYILVQNSEELQQQLGAIQDSQIFNETAVRLGQE NGYSFTADEVDAFLNEKIQQNNAELSDRELEAVAGGKSRCP IGTRFTFCVLVSSCWGSKC |
| *Mojavia pulchra JT2-VF2* | MSKQNVEQFYIVVENNQQLQEQLGQADNPQSFYDMAARL GQENGYSFTAQEAEDFTNQRVQQGNAELSDQNLEAVAGG RSCPMDTRFTFCVFISSCWGSKC |
| *Gloeotrichia echinulata HAB0833* | MSKENLEQFYILVQNSEALQQQLGAIQESEVFNETAARLGQ ENGYSFSAEEVDTFLNEKKQQSNAELSDRELEAVAGGKGE RCPINTRFTVCFLVSGCWGSRC |
| *Aliinostoc sp. HNIBRCY26* | MNNQNLYQFYTLIQNSQELQAQLGAAESPETFAETAVRLGE ENGYSFTTEDVNAFVSQQASRANADLSEADLEAVAGGKAH TCPIDTRFTFCVFISECWGSVC |
| *Nostoc sp.* | MSKENLEQFYIFVENNQQLQEQLGQAENPQSFYEKAVQLG QENGYSFTVQEAEDFISQKAQQGNAELSDQQLEAVAGGKG GCPLDTKFTFCLLISSCLGSKC |

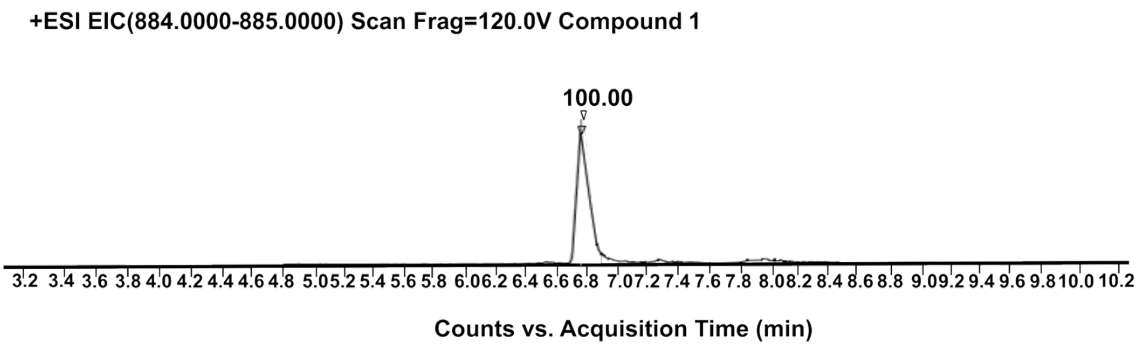


+ESI Scan (rt: 6.829 min) Frag=120.0V Compound 1

| MH$^{+1}$(mono) | MH$^{+2}$(mono) | MH$^{+3}$(mono) | MH$^{+4}$(mono) | MH$^{+5}$(mono) |
|---|---|---|---|---|
| 2651.2138 | 1326.1105 | 884.4095 | 663.5589 | 531.0486 |

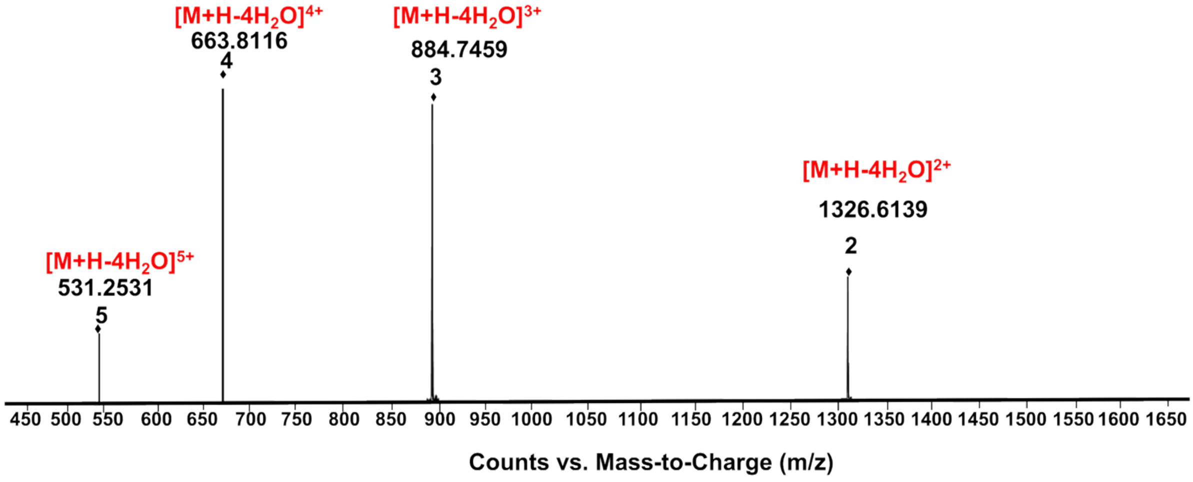


**Figure S1.** LC-QTOF-ESI/MS profile of nostolysamide C (positive mode) showing the monoisotopic mass for the different charge states for the 4-fold dehydrated product.

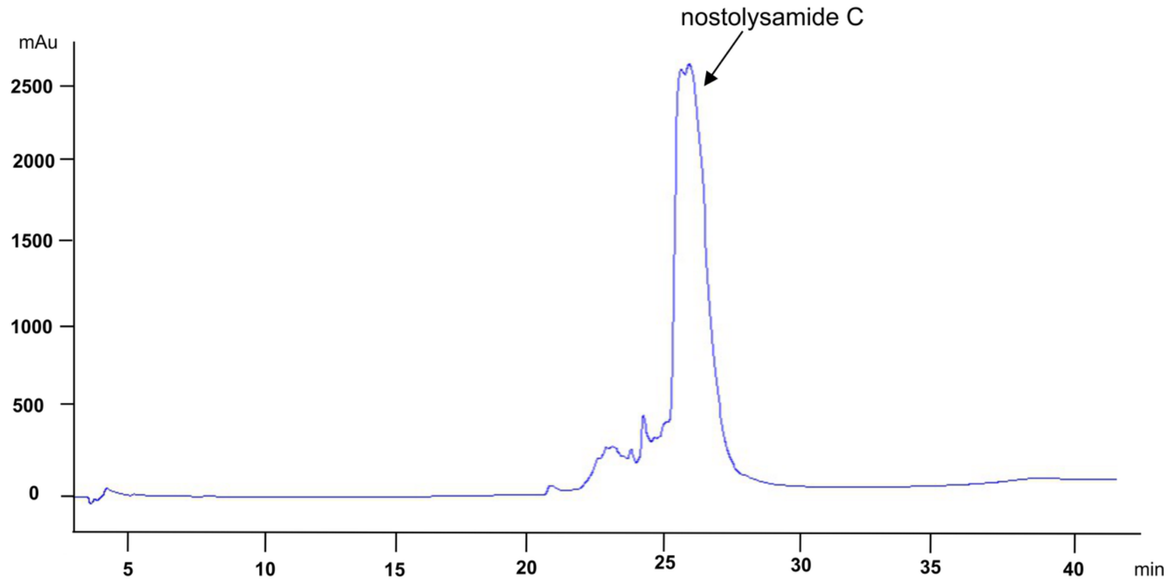


**Figure S2.** Reversed-phase HPLC trace of purified nostolysamide C (**1**), which elutes around 47% acetonitrile.

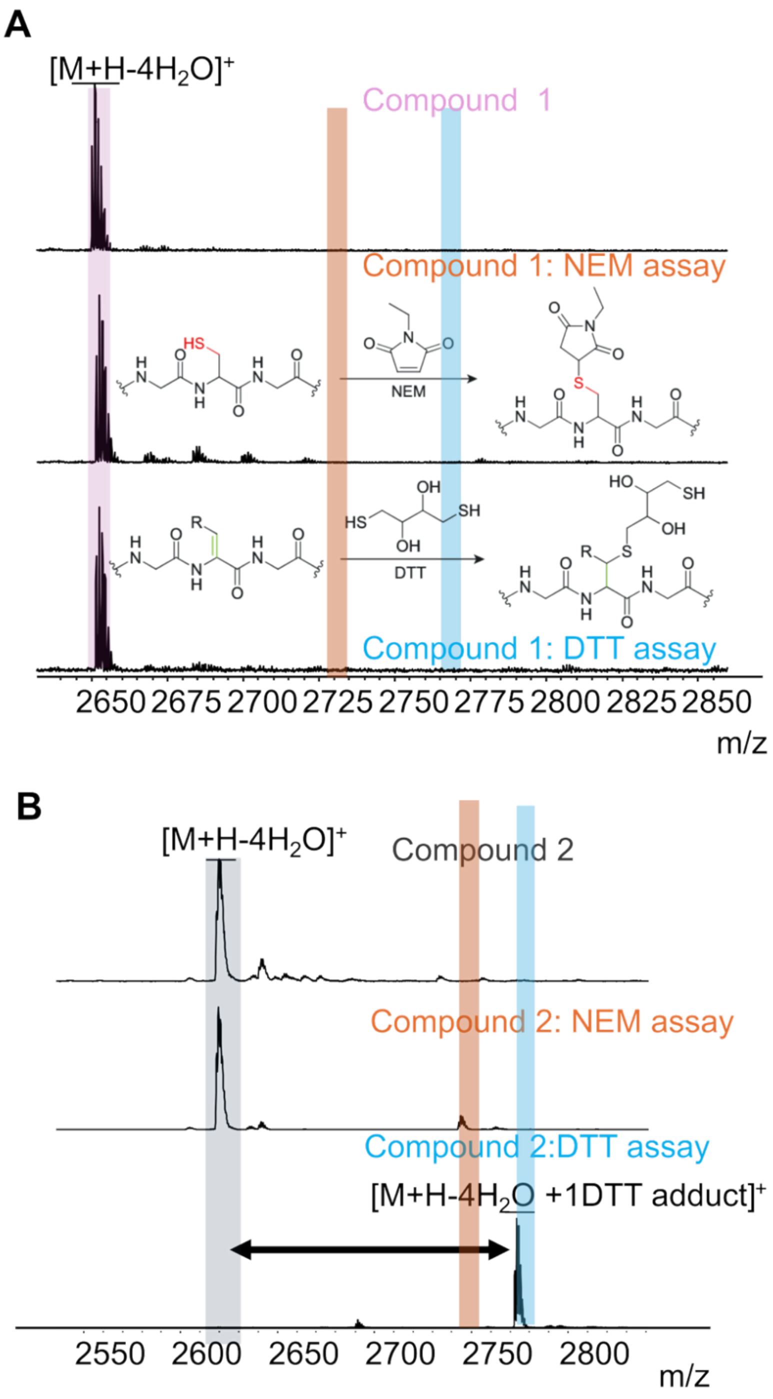


**Figure S3. MALDI-TOF mass spectra of NEM and DTT assays.**[1] (**A**) Compound **1** m/z: observed $[M+H-4H_2O]^+$=2651.3; calculated =2651.2. No adduct was observed when **1** was treated with NEM (expected m/z shown by red highlighted band) or DTT (expected m/z shown by blue highlighted band), indicating that the peptide is fully cyclized. (**B**) Compound **2** m/z: observed $[M+H-4H_2O]^+$= 2618.7; calculated= 2619.2. While no NEM adducts were formed showing the absence of a free Cys, a DTT adduct was observed, suggesting one dehydroamino acid in the product, as expected for this mutant. m/z observed $[M+H-4H_2O +1DTT\ adduct]^+$= 2772.5; calculated mass =2773.2.

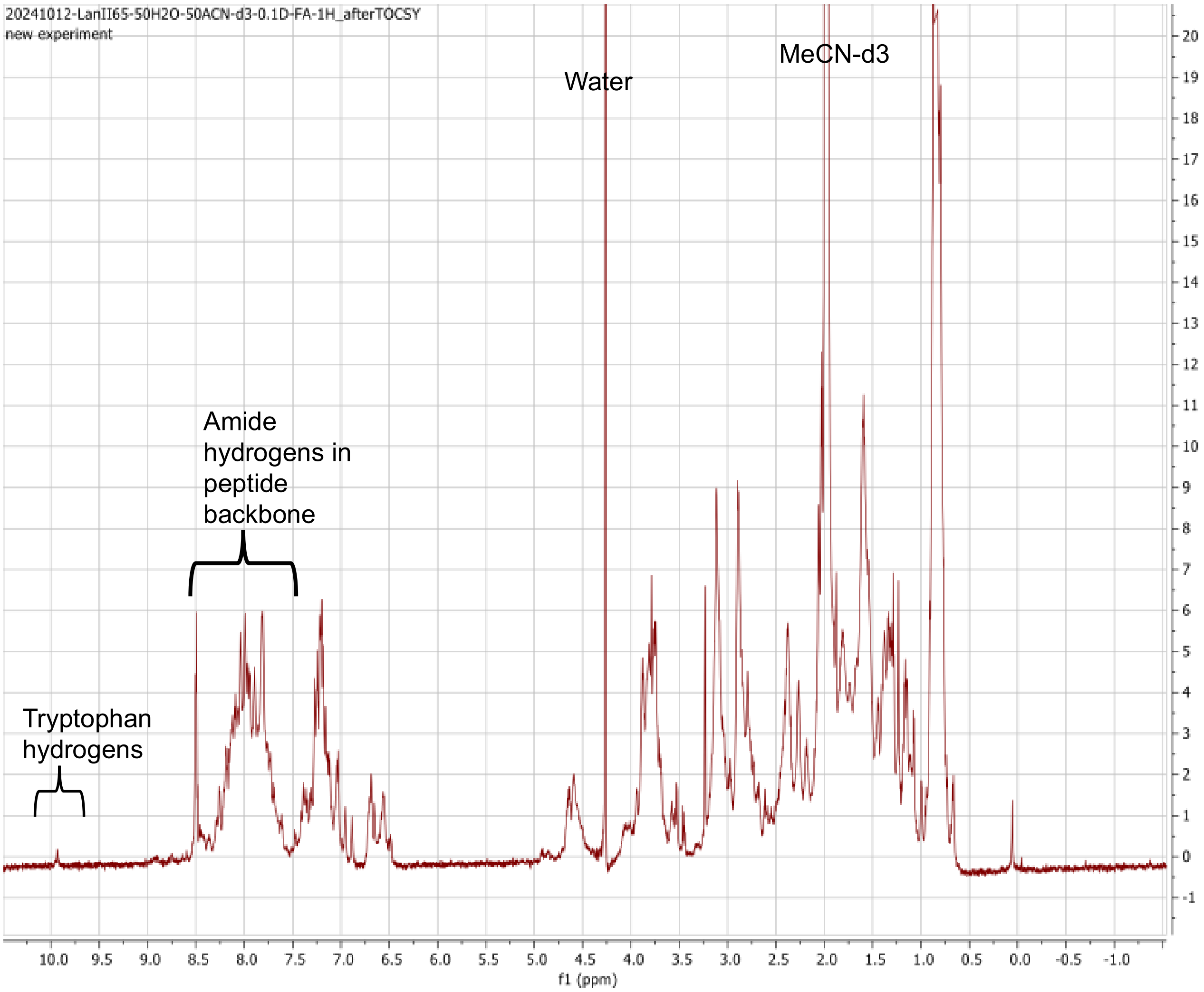


**Figure S4:** $^{1}H$ NMR spectrum of nostolysamide C (**1**) in 50% ACN-$d_3$/$H_2O$. The spectrum shows severe signal overlap in the amide region and poor signal dispersion. Furthermore, the solubility of the peptide was poor. Similar observations were made in other solvent systems.

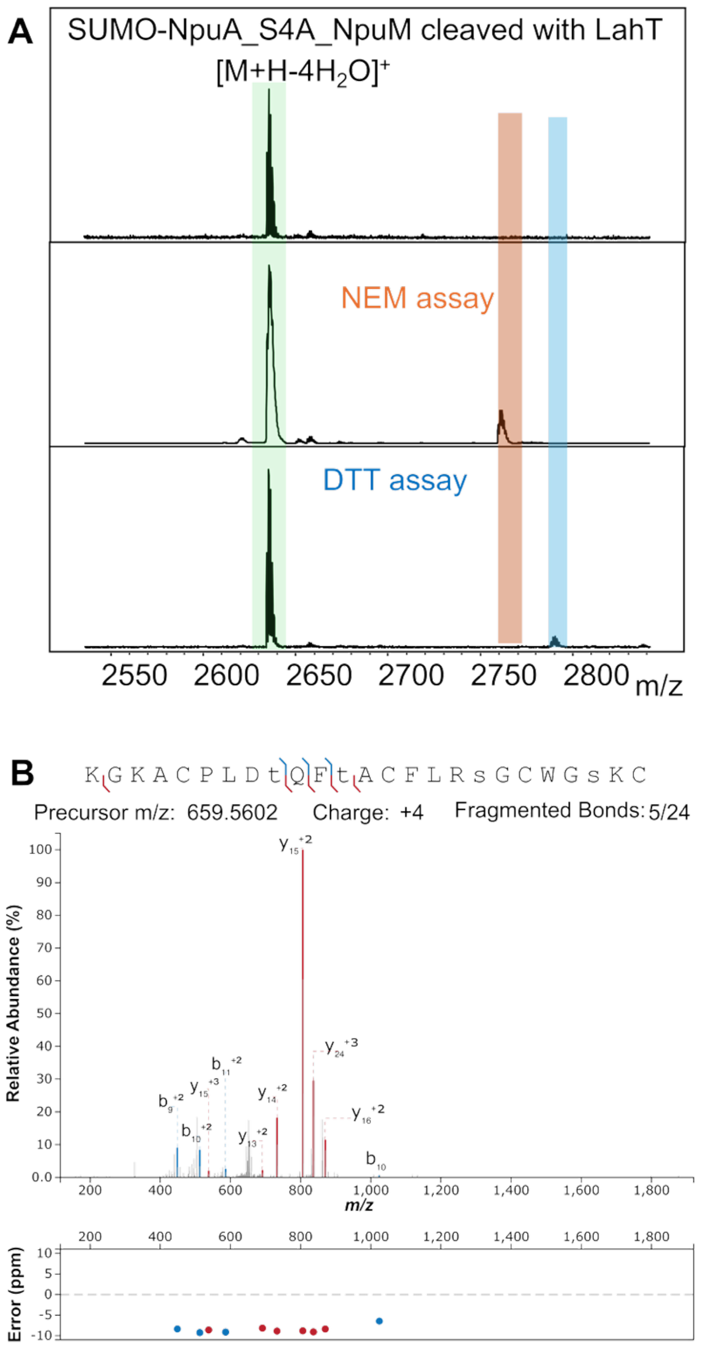


**Figure S5. Characterization of SUMO-NpuA-S4A modified by NpuM in *E. coli* and cleaved in vitro with LahT150.** (**A**) MALDI-TOF MS m/z observed $[M+H-4\ H_2O]^+=2634.3$; calculated $[M+H-4\ H_2O]^+=2635.2$. DTT and NEM assays showed no adducts, suggesting the serine at position 4 is not involved in the four dehydrations in WT NpuA. Furthermore, these assays show that the mutant peptide is fully dehydrated and mostly cyclized. (**B**) HR-MS of SUMO-NpuA-S4A modified by NpuM and cleaved with LahT150 showing b and y ions when the input sequence (lower-case letters) involved dehydrated residues at the indicated positions. Figure made using IPSA.[2] The fragmentation pattern is similar to wild-type NpuM-modified NpuA (**Fig. 2B**). HR-MS m/z calculated $[M+H-4\ H_2O]^{4+}=659.5602$ and observed $[M+H-4\ H_2O]^{4+}=659.5602$.

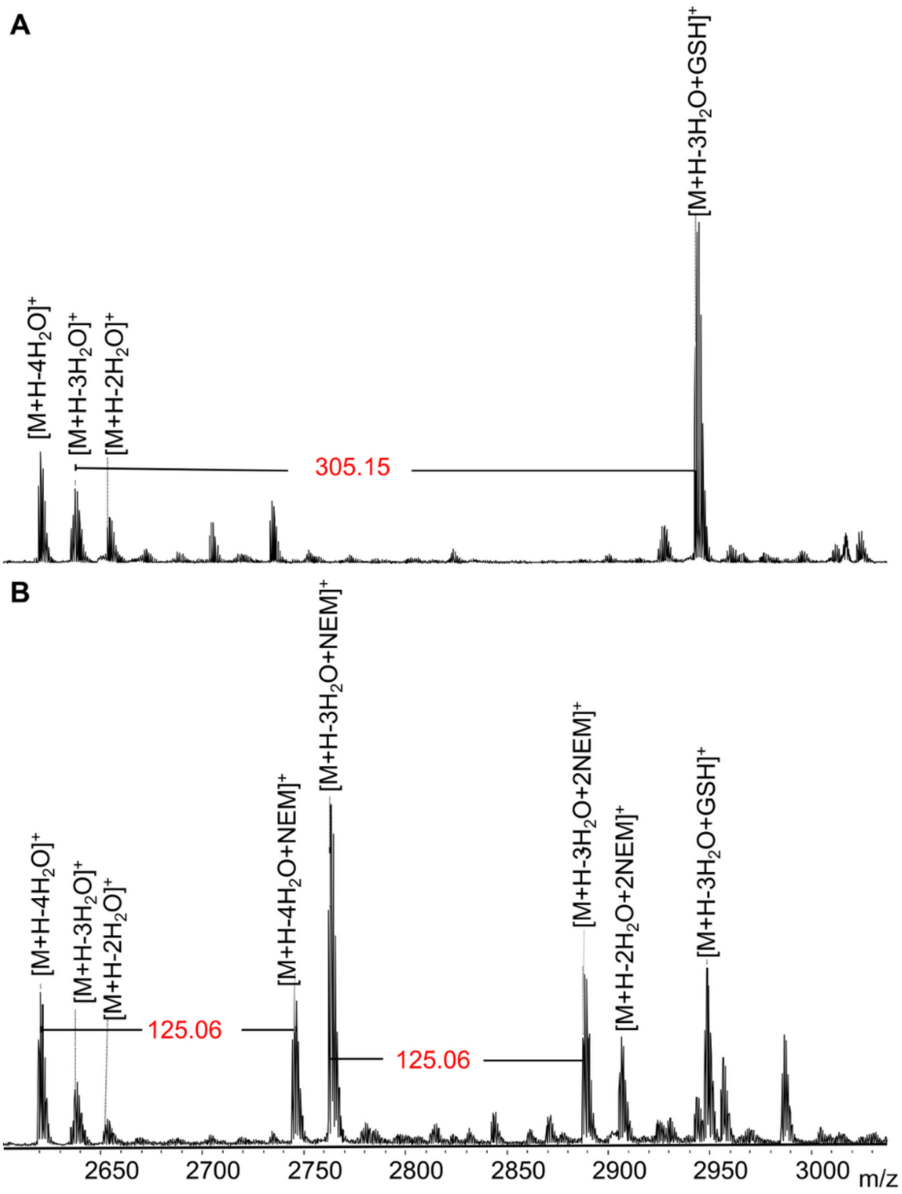


**Figure S6.** MALDI-TOF mass spectra of (A) NpuA-C14A co-expressed with NpuM, and cleaved with LahT150. MALDI-TOF MS showed a mixture of dehydrations, m/z observed $[M+H-4\ H_2O]^+$=2619.2, calculated $[M+H-4\ H_2O]^+$=2619.2; m/z observed $[M+H-3\ H_2O]^+$=2636.6, calculated $[M+H-3\ H_2O]^+$= 2637.2; m/z observed $[M+H-2\ H_2O]^+$=2653.5, calculated $[M+H-2\ H_2O]^+$= 2655.3 (loss of -2 Da likely because of disulfide bond formation). A glutathione adduct was observed on the three dehydrated products, resulting from a dehydroamino acid that was not cyclized with its partner Cys. (B) NEM assay showed several NEM adducts, which resulted in a +125 Da increase in m/z.

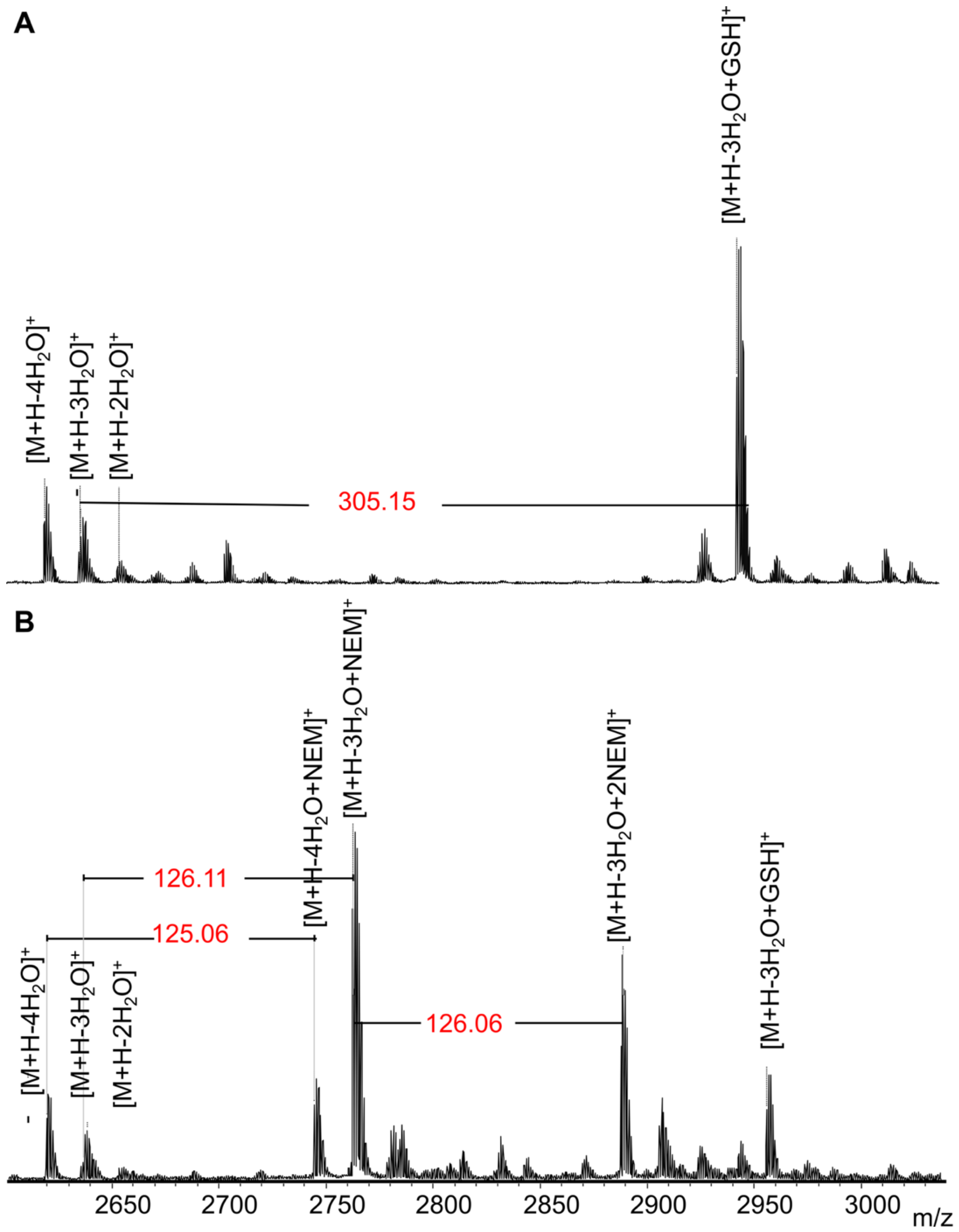


**Figure S7.** MALDI-TOF mass spectra of (A) NpuA-C20A co-expressed with NpuM, and cleaved with LahT150. MALDI-TOF MS showed a mixture of dehydrations, m/z observed $[M+H-4\ H_2O]^+$=2619.5, calculated $[M+H-4\ H_2O]^+$=2619.2; m/z observed $[M+H-3\ H_2O]^+$=2636.5, calculated $[M+H-3\ H_2O]^+$= 2637.2; m/z observed $[M+H-2\ H_2O]^+$=2653.5, calculated $[M+H-2\ H_2O]^+$= 2655.3 (loss of -2 Da likely because of disulfide bond formation). A glutathione adduct was observed on the three dehydrated products, resulting from a dehydroamino acid not being cyclized with its normal partner Cys. (B) NEM assay showed several NEM adducts, which resulted in a +125 Da increase.

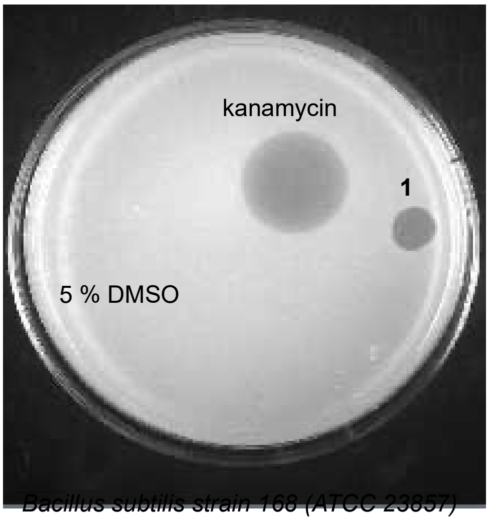


**Figure S8. Bioactivity assay with nostolysamide C (1) against *B. subtilis*.** A zone of bacterial growth inhibition was observed with *B. subtilis* 168 for both the positive control kanamycin (1 µL, 86 mM) and the tested compound **1** (2 µL,100 µM), while 5% DMSO (2 µL) served as a negative control.

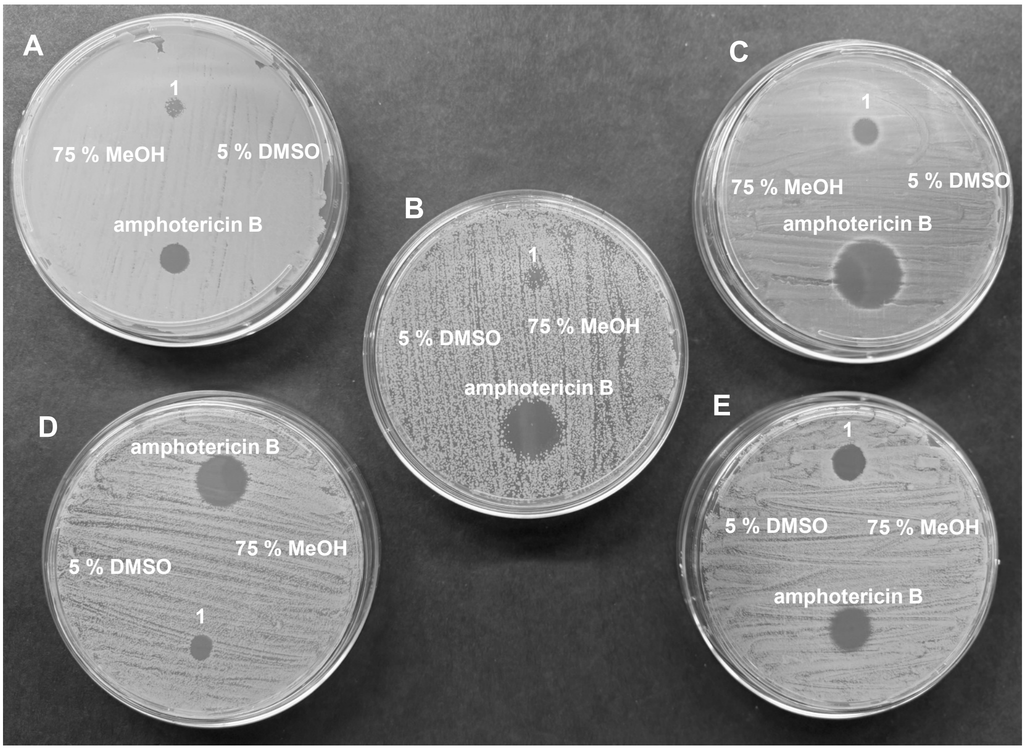


**Figure S9. Bioactivity assay of nostolysamide C (1) against *Candida* species**. Zones of growth inhibition were observed for compound **1** (2 µL, 400 µM) with *(***A**) *Candida krusei,* (**B**) *Candida albicans SN250*, (**C**) *Candida glabrata*, (**D**) *Candida albicans*, and (**E**) *Candida tropicalis.* Amphotericin B (2 µL, 1 mM) was used as the positive control and 5% DMSO (2 µL) and 75% MeOH (2 µL) served as negative controls.

**A**

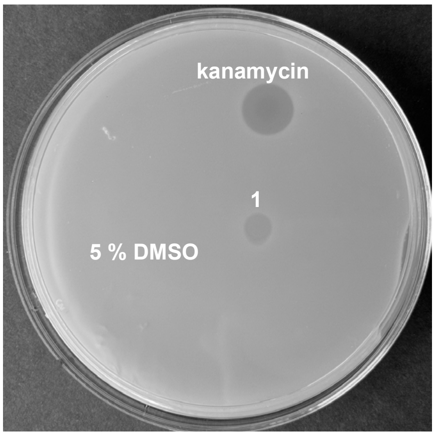


*Escherichia coli*

**B**

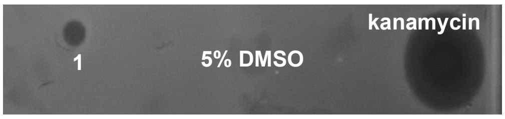


*Klebsiella pneumoniae ATCC 27736*

**Figure S10. Bioactivity assay with nostolysamide C (1) against gram–negative bacteria.** (**A**, **B**) Zones of bacterial growth inhibition were observed with *E. coli* (**A**) and *K. pneumoniae* (**B**) for the positive control kanamycin (2 µL, 0.86 mM) and compound **1** (2 µL, 200 µM); 5% DMSO (2 µL) was used as negative control.

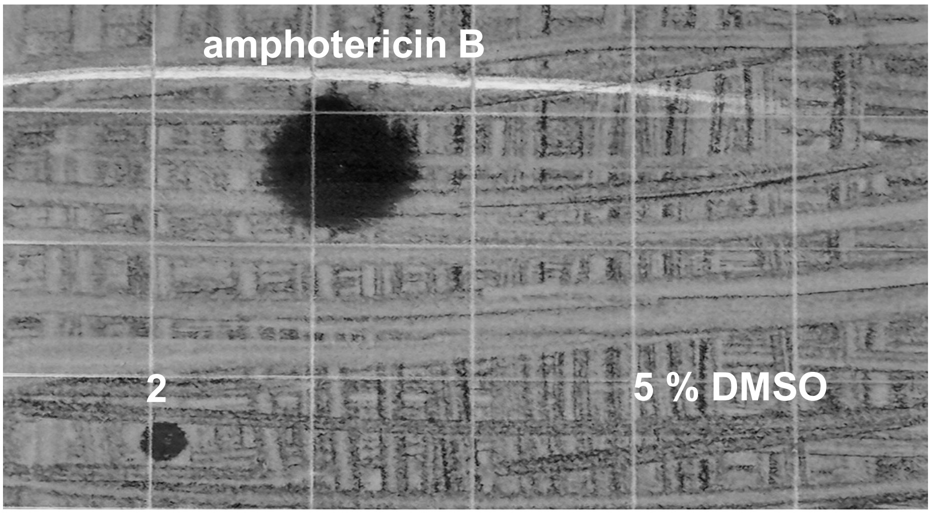


**Figure S11. Bioactivity assay with nostolysamide C-C25A against *Candida tropicalis*.** Zones of bacterial growth inhibition were observed against *C. tropicalis* with the positive control amphotericin B (2 µL, 1 mM) and compound **2** (2 µL, 200 µM); 5% DMSO (2 µL) served as the negative control.

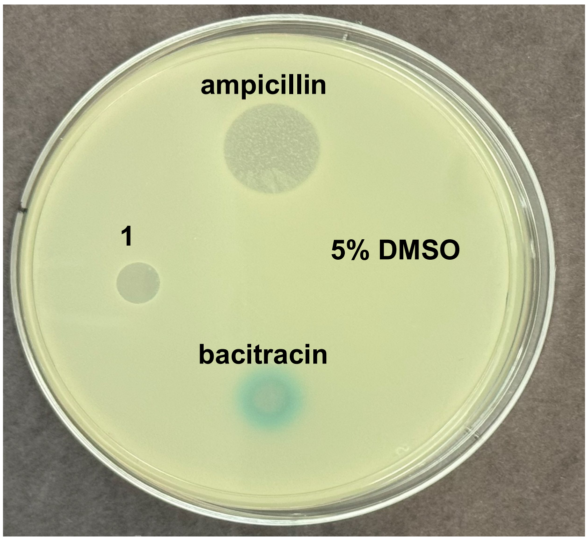


*Bacillus subtilis BSF2470 (1A980)*

**Figure S12**. To assess whether compound **1** targets lipid II, a reporter strain of *B. subtilis* was used with LacZ expression.[3] The soft agar contained X-gal. Amounts of compounds spotted were: 2 µL of 100 µM of compound **1**, 2 µL of 35 mM solution of the positive control bacitracin, and 2 µL of a 269 mM solution of the negative control ampicillin. As another control, 2 µL of 5% DMSO was also spotted.

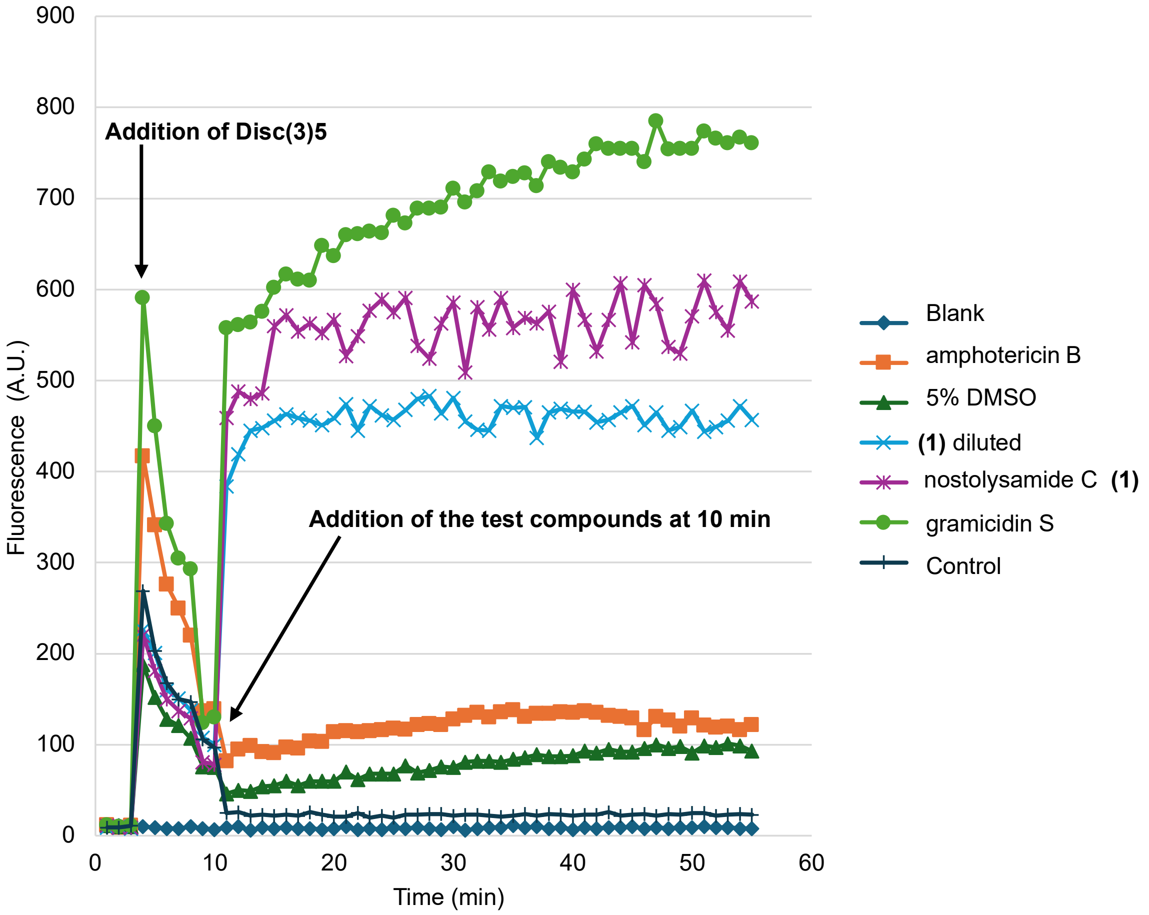


**Figure S13. Membrane depolarization measured by DiSC$_3$(5) fluorescence.** Fungal membrane potential was assessed using the voltage-sensitive dye DiSC$_3$(5). *Candida tropicalis* cells were incubated with DiSC$_3$(5) until fluorescence stabilized, followed by treatment with various test compounds/antibiotics. Increased fluorescence indicates dye release due to membrane depolarization. Gramicidin S (10 µL, 1 mM stock, 1 mM final) was used as the positive control. Nostolysamide C (**1**) (10 µL, 50-100 µM stock) showed similar fluorescence traces over time, suggesting nostolysamide C causes membrane depolarization.

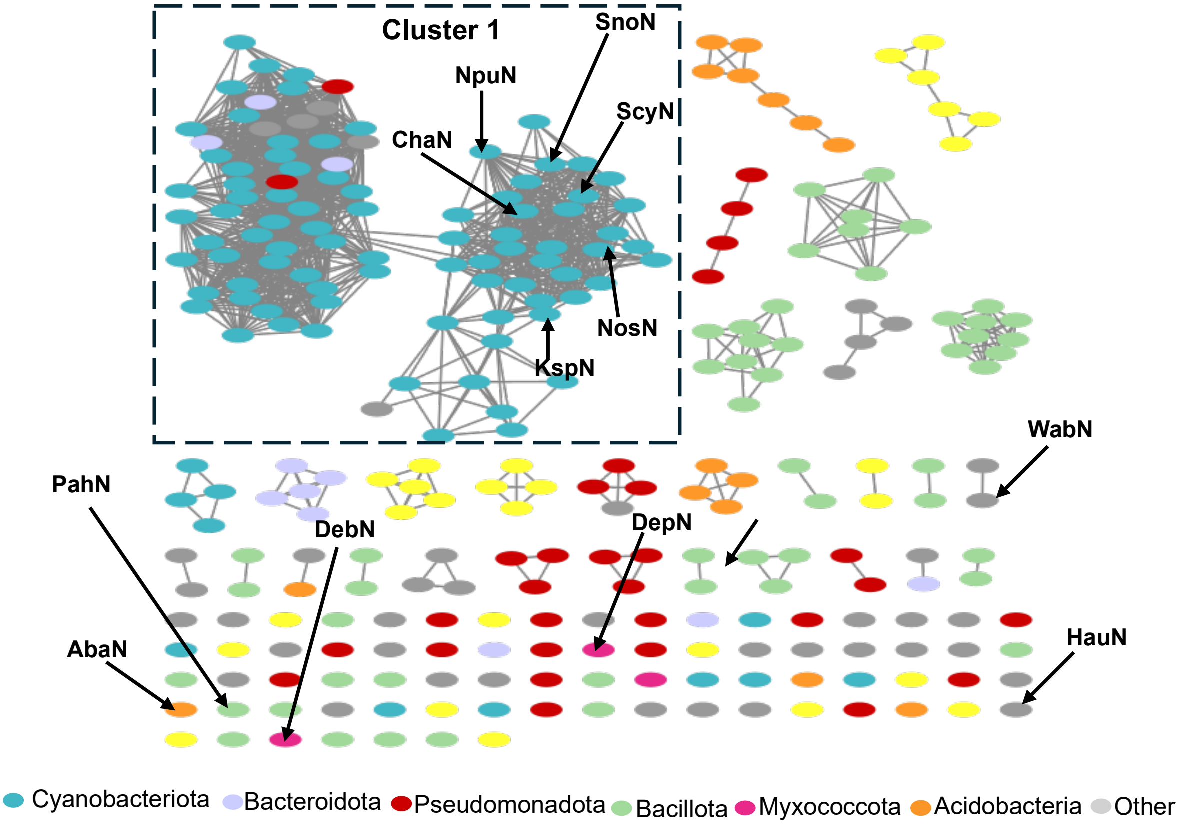


**Figure S14**. Sequence similarity network (SSN) of a subset of the GNAT-family acetyltransferases colored by the phyla they are encoded in. Protein sequences were obtained from UniProt via a BLAST search using NpuN as the query. The network was created by linking sequences sharing ≥ 70% pairwise identity. Nodes represent individual sequences, while edges indicate sequence similarity above this threshold. SSN construction and visualization were performed with the EFI-EST web tool with default parameters (minimum length 0, maximum length 50,000, E-value filter ≤100).[8] NpuN_repnode-1.00 is visualized in Cytoscape v3.10.[9] Highlighted is cluster 1, which includes NpuN and other multidomain acetyltransferases. Also indicated are other acetyltransferases identified by Piel and coworkers that are likely involved in RiPP biosynthesis.[10]

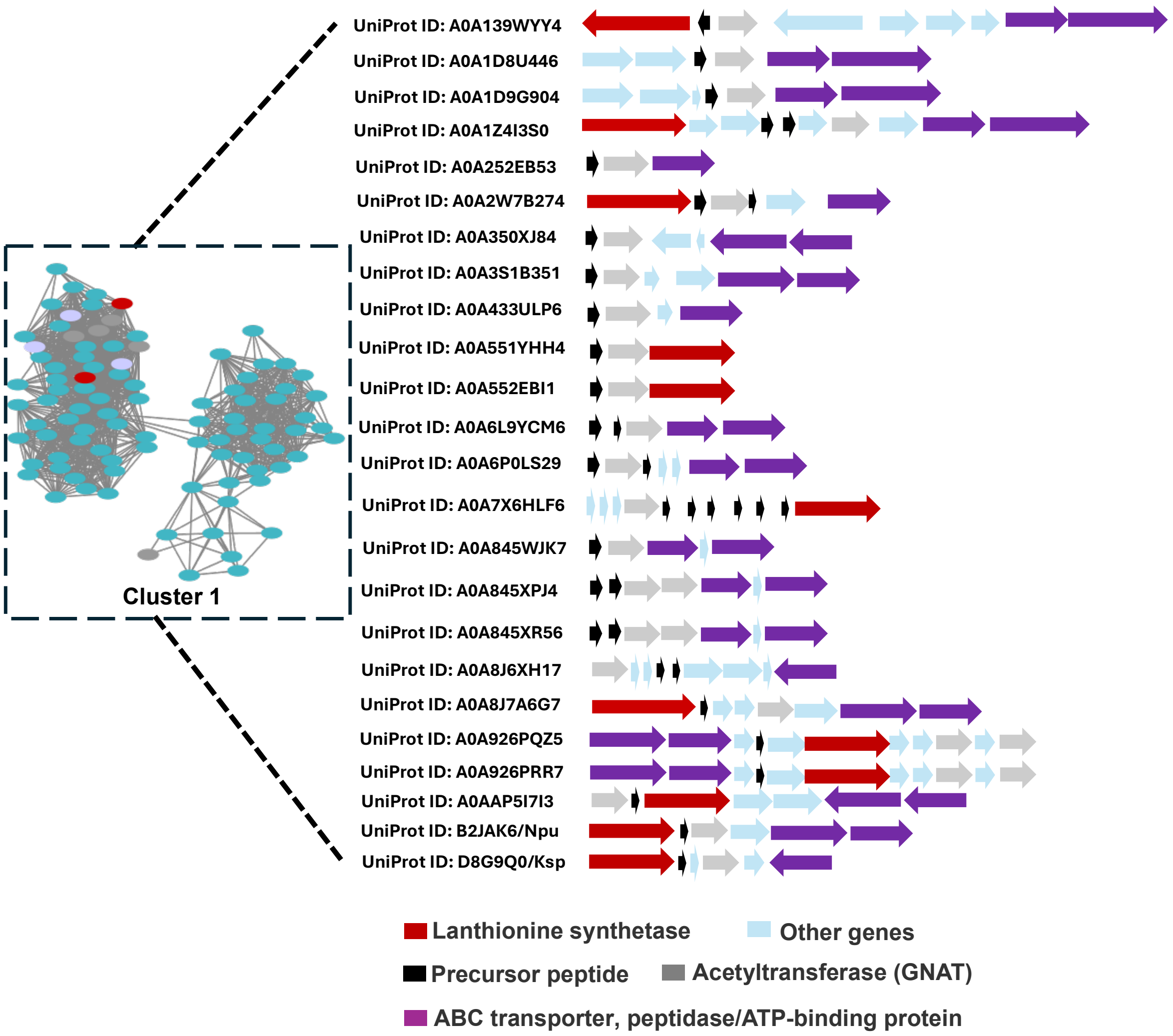


**Figure S15**. A representative subset of genome neighborhoods from Cluster 1 was identified using the Enzyme Function Initiative–Genome Neighborhood Tool (EFI-GNT). These neighborhoods were filtered to include BGCs encoding a conserved precursor peptide gene along with an acetyltransferase and/or lanthionine synthase. Coloring in the SSN is by phyla as in Fig. S14. Coloring of the BGCs is as shown in the legend of the Figure.

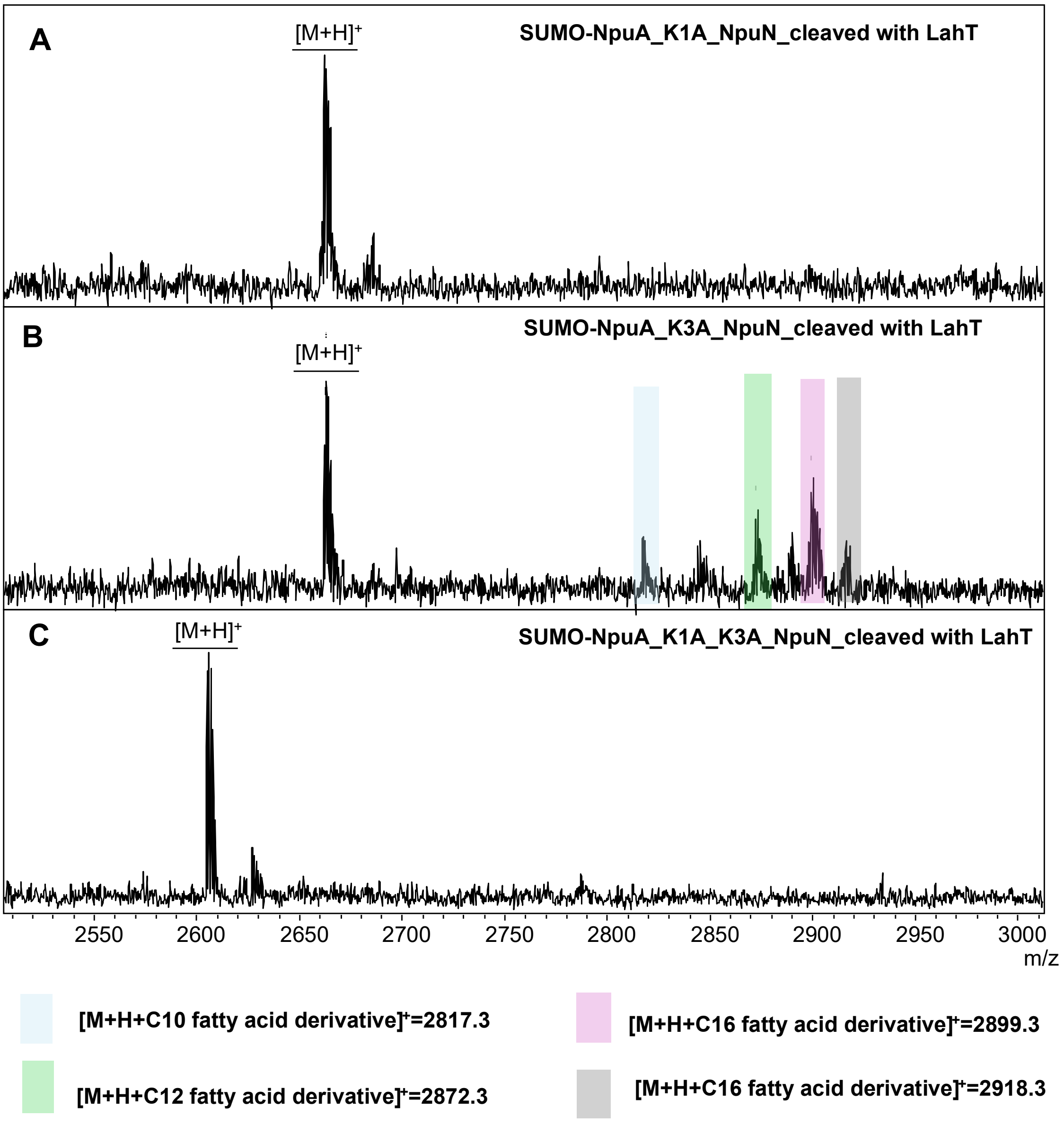


**Figure S16. MALDI-TOF MS of the site-directed mutagenesis products of the lysine residues in the core peptide of NpuA to identify where acylation occurs.** (**A**) SUMO-NpuA-K1A co-expressed with NpuN after cleavage with LahT150, m/z observed $[M+H]^+$= 2662.1, calculated 2666.2. No acylation was observed. (**B**) Lysine at position 3 of the core peptide of SUMO-NpuA was mutated to alanine and co-expressed with NpuN, resulting in products with different acyl groups as shown by MALDI-TOF MS. (**C**) Double mutation of the lysine residues at positions 1 and 3 of the core peptide of NpuA to alanine. No lipidation was observed.

A

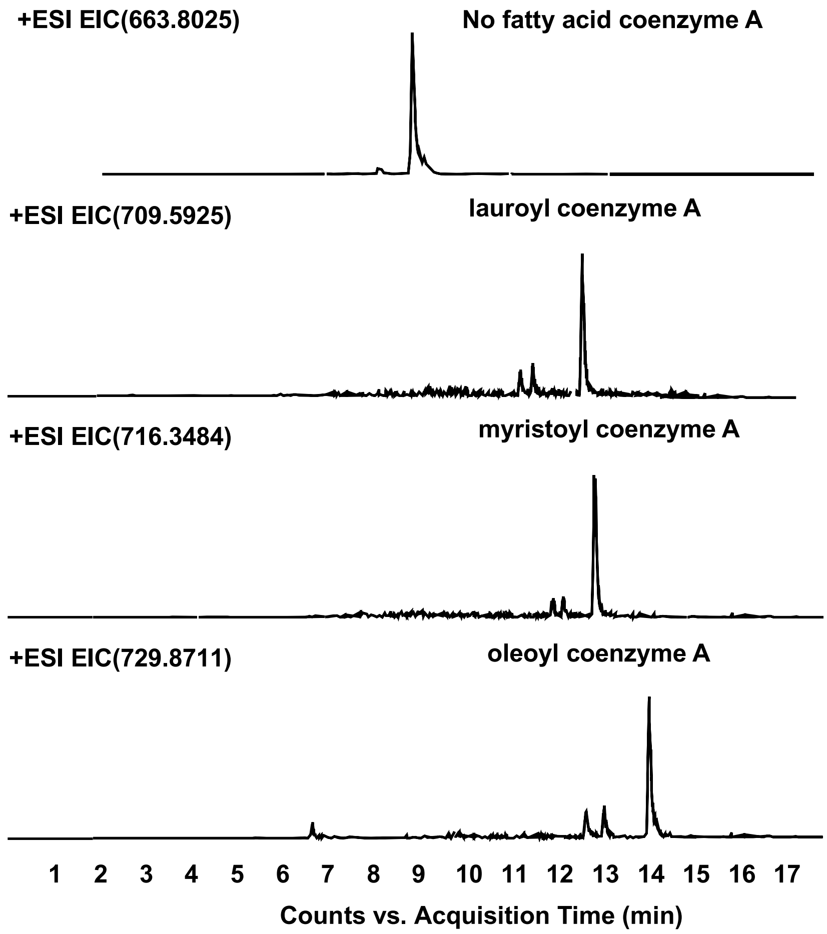


B

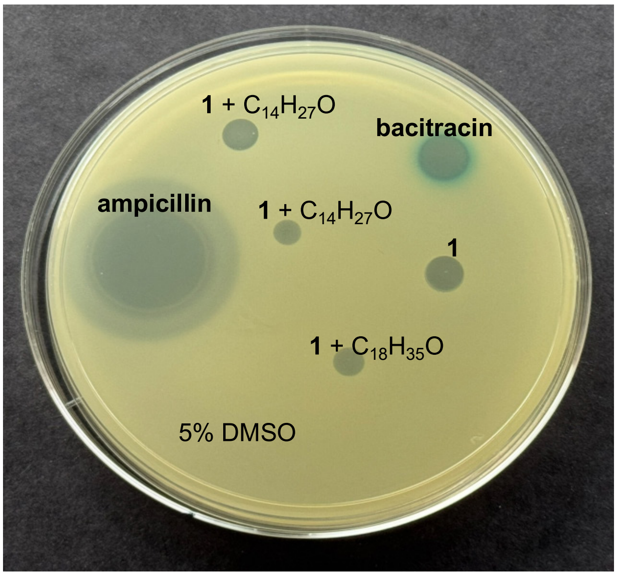


*Bacillus subtilis BSF2470 (1A980)*

**Figure S17.** (**A**). An agar diffusion assay was used to assess whether nostolysamide C and acylated variations target lipid II, using a reporter strain of *B. subtilis* with LacZ expression. The soft agar contained X-gal. Spotted were 2 μL of 100 μM of compound **1** and 2 μL of the same solution treated with NpuN and the indicated acyl-CoA (MS spectra shown in panel C).

Also spotted were 2 µL of a 35 mM solution of the positive control, bacitracin, and 2 µL of a 269 mM solution of the negative control, ampicillin. As another control, 2 µL of 5% DMSO was also spotted. (**B**) Extracted ion chromatograms (EICs) showing electrospray ionization MS analysis of the in vitro reactions of His-SUMO-NpuN (20 µL, 83 µM), nostolysamide C (20 µL, 250 µM, and different acyl-CoAs (10 µL, 1 mM).

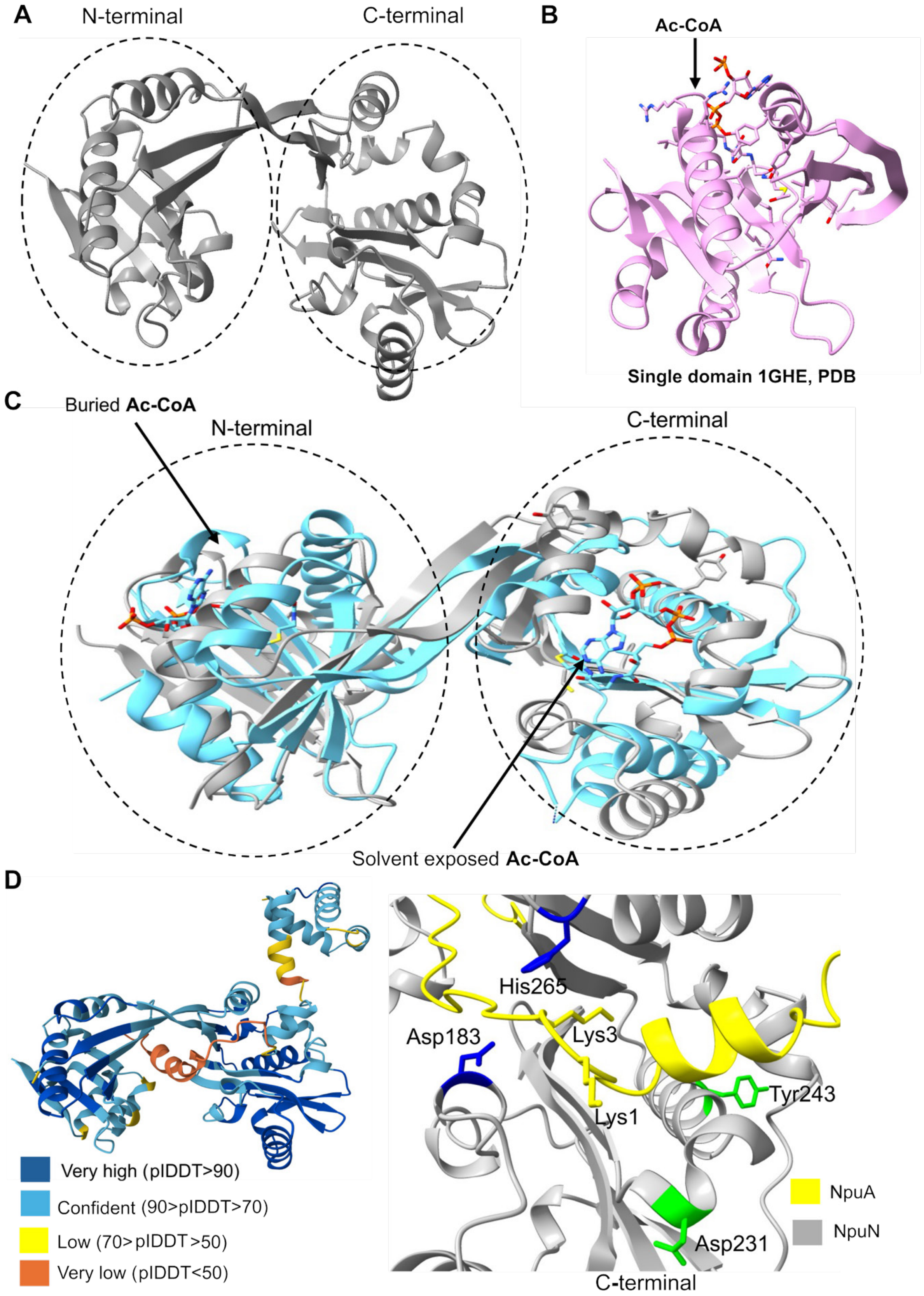


**Figure S18. Structural comparison of characterized and AlphaFold-predicted[4] GNAT family proteins with single and tandem domain architectures.** (**A**) AlphaFold3 model of NpuN, the tandem-domain GNAT enzyme investigated in this study, featuring two acetyltransferase-like domains connected by a linker region. (**B**) The single-domain GNAT tabtoxin resistance (TTR) protein obtained from a DALI search displays the typical α/β acetyltransferase fold of the Gcn5-related family, with Ac-CoA docked (PDB ID 1GHE). (**C**) Structural alignment of two tandem GNAT proteins: grey for NpuN; blue for Virulence-induced protein F (VipF) from *Legionella hackeliae* with acetyl-CoA bound (PDB ID 6WQB).[5] The alignment illustrates how Ac-CoA might bind to the N- and C-terminal domains of NpuN, with one Ac-CoA buried vs one solvent exposed. (**D**) AlphaFold3 model of NpuN with the substrate NpuA colored by predicted Local Distance Difference Test (pLDDT) values and visualized using UCSF ChimeraX (left).[6] A close-up of the predicted C-terminal active site of NpuN with the substrate NpuA is shown on the right. Two residues, Asp183 and His265, in the C-terminal GNAT domain are positioned within approximately 5 Å of the substrate Lys1 ε-amino group that is acylated by NpuN, suggesting possible roles in acyl transfer. Conversely, a Tyr that has been implicated as general acid to protonate the thiolate leaving group in many GNATs[7], including VipF,[5] is conserved in NpuN (Fig. S19). However, this Tyr243 is not close to Lys1 of NpuA in the model of NpuN-NpuA. Thus, location of acyl-CoA and NpuA binding and the catalytic residues cannot be confidently predicted by the AlphaFold model.

**A**

```
NpuN    ---------------MYTL-------NILKDSTA--VMYEKMTFPAFQY-----------  25
TTR     MNHAQLRRVTAESFAHYRHGLAQLLFETVHGGASVGFMADLDMQQAYAWCDGLKADIAAG  60
                        *         : ::..::  .* :     *: :

NpuN    --VLQEINSKE-SLIAIAASYQEEPVGLVLAEIRTYNQSAEVLSLFVEPSHRCAGIGTNL  82
TTR     SLLLWVVAEDDNVLASAQLSLCQKPNGL---------NRAEVQKLMVLPSARGRGLGRQL  111
          :*  :  ..:   * :    *  ::* **          : *** .*:* ** *  *:* :*

NpuN    LNRLETELTYRRCKKVELVYMTGKSTTPALENLLHKCGWTSPQPRMLVCKGEMEKGIQAP  142
TTR     MDEVEQVAVKHKRGLLHLDTEAGSVA----EAFYSALAYTR-------------------  148
        ::.:*   . ::   :.*   :*. :    * :    .:*

NpuN    WLKQYSQLPAAYTIFPWREITTQERQAIERQQATQPWIPEDLIPFKYDKQIEPLNSLGLR  202
TTR     ------------------------------------------------------------  148

NpuN    YQGQVVGWLIS--HRIFPDTIRYTCAFCRPDIQKKGRMITLFYEAAKLQLNAKIPHFIWT  260
TTR     -VGELPGYCATPDGRLHPTAIYF-KTLGQPT-----------------------------  177
          *:: *:  :   *:.* :* :  :: :*

NpuN    TPMNHQSMVAFVKYRCLPYLTSIQETRIISKLLLG   295
TTR     -----------------------------------   177
```

**B**

```
VipF    QGMIICNNQIDDDQLKAIQDLASLCREHDGGTPTFYNHLLIQKRPTENNVLYF---QDNQ  57
NpuN    ---MYTLNILKDSTAVMY--------EK-MTFPAF-QY-VLQEINSKESLIAIAASYQEE  46
NosN    ---MYELNILDNSTASAY--------EW-MTFPSY-KY-LLHTLDTDNTIVAIGATNNQQ  46
           :   * :.:.              *     *:: :: :::   :.:.:: :    :::

VipF    LLGFL--SVYFFYEDACEVSLIVSPLHRRQGIAKQLLQTIMPLLTAKEMTTLIFSTPTEI  115
NpuN    PVGLVLAEIRTYNQSAEVLSLFVEPSHRCAGIGTNLLNRLETELTYRRCKKVELVYMTGK  106
NosN    PVGLVLAQMVQ-DRIVVIFSIFVAIEERNKGIGTALLTRIQAELQDRGCKILQLNYTTGK  105
         :*::  .:    . .  .*::*   .*  **.. **  :   *  :  . : :   *

VipF    NDDWLINQGFSYRNSEYHMQRNGYDPIFMPTPKLHIRKATEDDIPALCAIDEACFPEHQE  175
NpuN    STT---------PALENLLHKCGWTS---PQPRMLVCKGEME------------KGIQA  141
NosN    PTT---------AALERVIEKCNWTP---PETRTLVCKGEAK------------RIIEA  140
                       *  :.: .:    *  :  : *.  .                :

VipF    NMISRFSMLLNDASYTLFLASY------NHIIVGKAHIHWQSKEAIF---SDIAIFRQYQ  226
NpuN    PWLKQYSQLP--AAYTIFPWREITTQERQAIERQQATQPWIPEDLIPFKYDKQIEPLNSL  199
NosN    NWIRRYSRLP--SSYSIFPWEEITPEERQIIQQQQTQPWIPEDLVPFKYEDNLEPLNSI  198
          : ::* *   ::*::*         : *   :    *  :: :     ..      :

VipF    GQGWGGELLSYCINQALTSGKNKLML-----DVETSNQNALHLYT---RLGFKTANVSDY  278
NpuN    GLRYQGQVVGWLISHRIFPDTIRYTCAFCRPDIQKKG-RMITLFYEAAKLQLNAKIPHFI  258
NosN    GLRYQGQVVGWLINHRIAPHTIRYTCSFVREDLQKMG-RIIILYAEACERQIKANIPNAI  257
        *  : *:::.: *.: :   . :        *::.  . . : *:     .  :::

VipF    WVIPLPQLLT------NWALE--------------- 293
NpuN    WTTPMNHQSMVAFVKYRCLPYLTSIQETRIISKLLLG 295
NosN    WTVPSFHQSMVSFVKNHWLPYLNAVEESRGTLKLI-- 292
        *. *   :          .
```

**Figure S19**. (**A**) Sequence alignment of the N-terminal domain of NpuN and TTR. The two proteins display 18.83% sequence identity. Highlighted in the pink box is the P-loop motif

Q/R-X-X-G-X-G/A, which anchors acetyl-CoA in TTR and VipF. This sequence motif was observed in NpuN at the N-terminus. (**B**) Multiple sequence alignment of tandem GNAT protein VipF with NpuN and NosN (see also Fig. S18). The P-loop motif is boxed solid in the N-terminal domains, and dashed in the C-terminal domains. Unlike VipF that has two canonical P-loops, NpuN and NosN have atypical P-loop motifs at the C-termini with an Asn replacing the Gln and an Arg replacing the second Gly. Two conserved residues in NpuN and NosN that are close to Lys1 of NpuA in an AlphaFold3 model of the complex (Fig. S18D) are highlighted in blue. Two other residues that have been implicated in catalysis in VipF[5] are highlighted in green. The sequence identity between VipF and NpuN is 25.1%, whereas NpuN and NosN have a sequence identity of 55.3%.

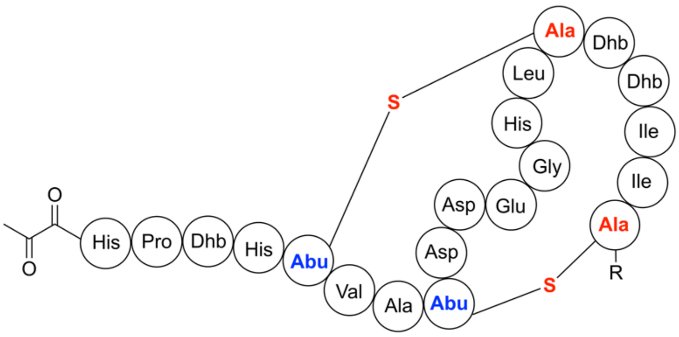


Pinensin A, R = Ala-OH
Pinensin B, R = OH

**Figure S20.** Chemical structures of pinensins A and B.[11]

```
NpuM   870  KFDIIGGAAGAILGLLALAQEQPMVLKQAIACGQHLLKYCNE—VFGKTSNL

ProcM  807  QLDLIGGCAGLIGSLVQLGTES—ALQWALRAGDHLIAQQNEEGAWSSSSSQ


NpuM   920  KGLIGFSHGAAGIAYALLRLYAVTQDTDYLDAARMAIAYENALFYPSVGNWR

ProcM  857  PALLGFSHGVAGYAAALAHLHAFSGDERYRIAAAAALAYERARFNKDAGNWP


NpuM   972  EITPVY--DPASLPVFWSTWCHGAPGIALGRLGSLSIDR-SEQILTDIEVA

ProcM  909  DYRSISRDSDSDEPSFMASWCHGAPGIALGRACLWGTALWDEECTKEIGIG


NpuM  1020  LKTTQNTALQTIDHLCCGNLG

ProcM  960  LQTTAAVSAVSTDHLCCGSLG
```

**Figure S21.** Sequence alignment of the C-termini of NpuM and ProcM. The Cys residues involved in $Zn^{2+}$ coordination in the active site are shown in red.
